# A feature-specific prediction error model explains dopaminergic heterogeneity

**DOI:** 10.1101/2022.02.28.482379

**Authors:** Rachel S. Lee, Yotam Sagiv, Ben Engelhard, Ilana B. Witten, Nathaniel D. Daw

## Abstract

The hypothesis that midbrain dopamine (DA) neurons broadcast an error for the prediction of reward (reward prediction error, RPE) is among the great successes of computational neuroscience^1–3^. However, recent results contradict a core aspect of this theory: that the neurons uniformly convey a scalar, global signal. For instance, when animals are placed in a high-dimensional environment, DA neurons in the ventral tegmental area (VTA) display substantial heterogeneity in the features to which they respond, while also having more consistent RPE-like responses at the time of reward^4^. We argue that the previously predominant family of extensions to the RPE model, which replicate the classic model in multiple parallel circuits, are ill-suited to explaining these and other results concerning DA heterogeneity within the VTA. Instead, we introduce a complementary “feature-specific RPE” model positing that DA neurons within VTA report individual RPEs for different elements of a population vector code for an animal’s state (moment-to-moment situation). To investigate this claim, we train a deep reinforcement learning model on a navigation and decision-making task and compare the feature-specific RPE derived from the network to population recordings from DA neurons during the same task. The model recapitulates key aspects of VTA DA neuron heterogeneity. Further, we show how our framework can be extended to explain patterns of heterogeneity in action responses reported among SNc DA neurons^5^. Thus, our work provides a path to reconcile new observations of DA neuron heterogeneity with classic ideas about RPE coding, while also providing a new perspective on how the brain performs reinforcement learning in high dimensional environments.

## Introduction

Among the more prominent hypotheses in computational neuroscience is that phasic responses from midbrain dopamine (DA) neurons report a reward prediction error (RPE) for learning to predict rewards and choose actions^1–3^. While clearly stylized, this account has impressive range, connecting neural substrates (the spiking of individual neurons and plasticity at target synapses, e.g. in striatum^6^) to behavior (trial-by-trial adjustments in choice tendencies^7^), all via interpretable computations over formally defined decision variables (**Fig. 1a**). The question of this article is whether and how the strengths of this account can be reconciled with a growing body of evidence challenging a core feature of the model, i.e. the identification of a scalar, globally broadcast RPE signal in DA responses.

**Figure 1:**
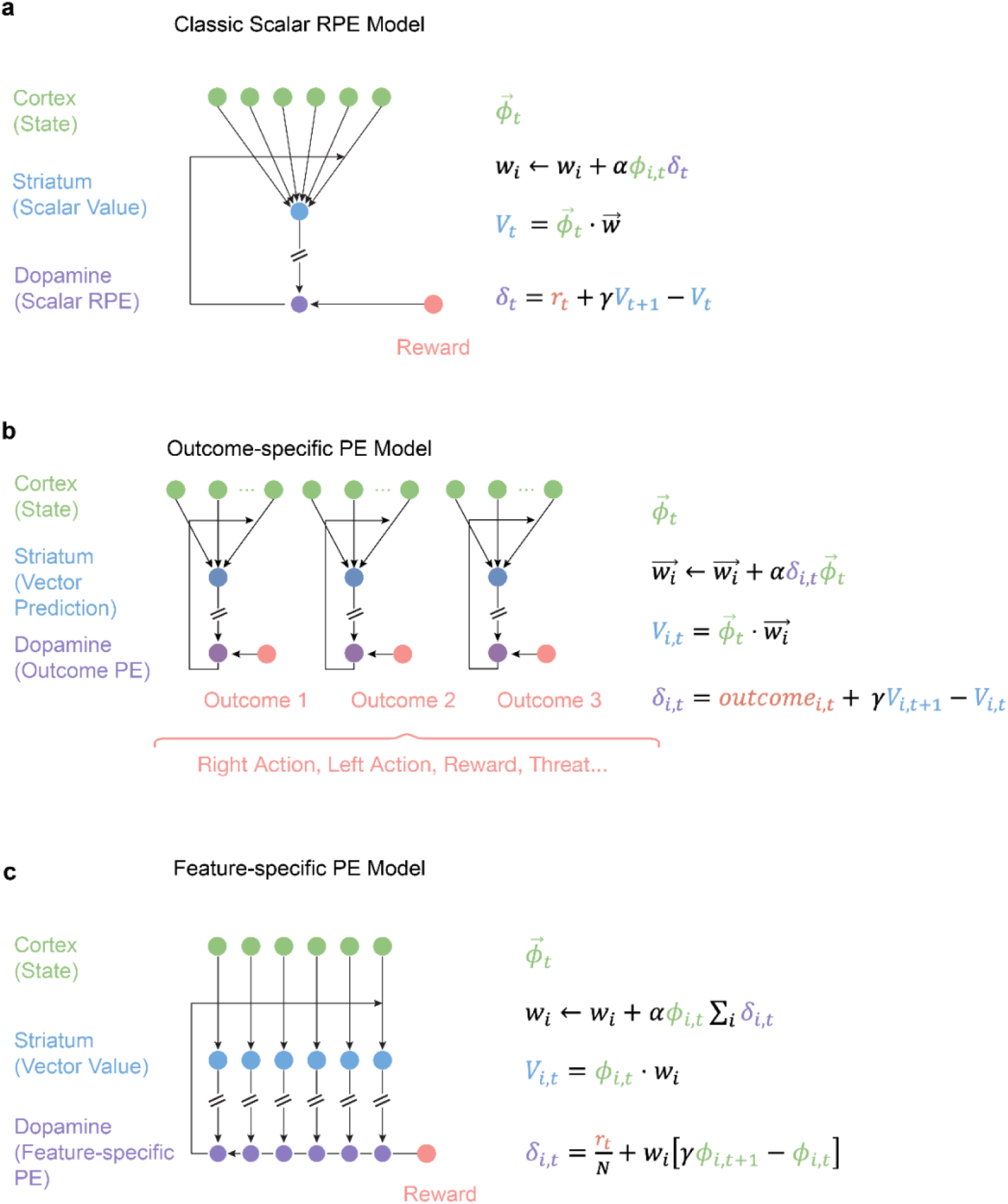
Feature-specific PE model updates classic TD learning to produce heterogeneous DA signals that reflect the state representation. **(a)** Classic mapping between equations of the TD learning model and brain circuitry^1–3^. **(b)** RL models can use different outcome variables to generate vector values and PEs. These outcomes can be driven by different aspects of reward^31,33,34^ (like in distributional RL models), specific actions (such as nosepoke, left, and right action)^40–42^, or other targets^35–39^. **(c)** Our proposed feature-specific PE model, which remaps the scalar RPE algorithm onto brain circuitry such that value and PE are vectors but the overall computations are preserved.

This scalar RPE is not a superficial claim of these theories, but instead one that connects a key computational idea to a number of empirical observations. Computationally, scalar decision variables reflect the ultimate role of any decision system in comparison: the decision-maker must order different outcomes against one another in ‘common currency’ to choose which to take ^8^. Anatomically, the ascending DA projection has historically been argued to have an organization more consistent with a scalar-like “broadcast” code (rather than a labeled line code): a relatively small number of individual neurons innervate a large area of the forebrain via diffuse projections^9–11^. Physiologically, early reports also stressed the homogeneity of responses of midbrain DA neurons on simple conditioning tasks – especially that a large majority of units respond to unexpected reward^12^.

However, the physiological argument for a scalar RPE is increasingly untenable, as a mounting body of recent work challenges the generality of this finding by demonstrating a range of variation in dopamine responses. Midbrain DA neurons can have heterogeneous and specialized responses to task variables during complex behavior^4,7,13–27^, even while often having relatively homogenous responses to reward^3,4,20,28,29^. In some areas, the signature response to reward can itself be blunted or reversed^5,7,15,21,30^.

How can we reconcile such DAergic heterogeneity with the substantial evidence for the RPE theory? There are a number of previous computational approaches relevant to this question, the vast majority of which share a common core hypothesis: that a set of different error signals could each serve to train different target functions, either predicting different aspects of reward ^31–34^, or predicting altogether different quantities^35–39^. For example, distributional RL suggests prediction errors (PEs) to predict different quantiles of reward expectation^33^, action prediction error models envision PEs to predict specific actions^40–42^, successor representation models suggest PEs for predicting different sensory stimuli^39^, and threat prediction error models suggest a PE to predict threat^34^. We refer to this class of models collectively as “outcome-specific PEs,” because they most often substitute a set of different prediction targets in place of the reward outcome in classic RPE (**Fig. 1b**).

Two implications of this class of explanations limit its applicability. First, the substitution of different targets in place of reward means these models cannot explain why most DA neurons respond to reward, nor how such uniformity can coexist with heterogeneity in other aspects of responding^4^. Second, anatomically, this approach amounts to duplicating the original scalar RPE model (**Fig. 1a**) for multiple types of error signals in parallel circuits (**Fig. 1b**). The fact that different targets need to be associated (at least roughly) in closed-loop fashion with their distinct PEs strongly suggests that DA neurons involved in different predictions must project to different target sites.

In contrast, here we propose a complementary new “feature-specific PE model” that offers an explanation for a distinct set of phenomena associated with heterogeneity *within* a projection-defined DA population, rather than across projections (**Fig. 1c**). In our new model, each DA neuron calculates a PE based on the subset of inputs it receives, with the full PE signal reconstructed at the target site (**Fig. 1c**). The key motivating observation is that within a single PE circuit (either the classic RPE, **Fig. 1a**, or any of its siblings in the outcome-specific models, **Fig. 1b**), the predictions being learned depend on an input variable known in RL models as “state”: a high-dimensional group of all sensory and internal variables relevant to the prediction. Since these are likely widely distributed throughout the brain, it is anatomically implausible that they converge uniformly and homogeneously on DA neurons to produce a scalar PE. Although the ascending DAergic projection is relatively diffuse^43^, inputs to DA neurons are not homogenous but instead arise from cortico-basal-ganglionic circuits that are highly topographically organized^43–49^. Here we propose that a distributed code for state is carried by corticostriatal circuits, which transform it into corresponding distributed codes for value and RPE that, in effect, decompose these scalar variables over state features^45^. In this way, different striatal and DAergic neurons reflect the contribution of different state features to value and RPE, but the ensemble collectively represents canonical RL computations over the scalar variables. In turn, this core circuit may be repeated in parallel between different targets, producing distinct aspects of heterogeneity between versus within projection-defined populations.

We initially investigate the feature-specific PE model by focusing on a recent study from our labs^4^ which provides one of the most detailed and dramatic examples of variability at the level of single neurons within the VTA, the nucleus classically most identified with an RPE. By performing 2- photon imaging while mice performed an evidence accumulation task in a virtual reality T-maze, we observed that while neurons respond relatively homogeneously to reward during the outcome period, during the navigation and decision period, they respond heterogeneously to kinematics, position, cues, and more^4^. We show how our feature-specific RPE model explains these data, while prominent outcome-specific models cannot. Moreover, we demonstrate that our model can be integrated with the outcome-specific PE framework to explain aspects of SNc DA neuron cell-to-cell heterogeneity^5^.

Our new model offers a number of key insights. First, the contrast between the feature-specific and outcome-specific models establishes a broad framework predicting that distinct outcome-vs feature-related aspects of DAergic heterogeneity predominate between-vs within projection-defined DA populations with different targets, which immediately suggests direct experimental tests. Second, within the canonical VTA RPE circuit, the feature-specific model explains a striking contrast observed in dopamine neurons: heterogeneous responses to task variables alongside much more uniform outcome-period responses^4,20^. This is a key empirical signature of the distributed value code we posit, but poorly explained by the earlier class of explanations. Third, the model retains an algebraic mapping to the standard theory^1–3^, preserving its successes while improving its match to anatomical and physiological evidence. Finally, the theory exposes an unexpected connection between the puzzling empirical phenomena of DAergic heterogeneity and a major theoretical question in RL models in neuroscience: the nature of state. While there has been substantial theoretical interest in the principles by which the brain represents task state^50–53^, there is relatively little empirical evidence to constrain these ideas. The new model suggests that the DA population representation itself can provide a window into this hitherto elusive concept of the neural representation of state.

## Results

### The Feature-Specific PE model

Here, we propose the feature-specific PE model as an extension of the classic scalar RPE model (**Fig. 1a**). RL models typically assume that the goal of the learner is to learn the value function *V*(*s*_*t*_) = *E*[*r*_*t*_ + *γr*_*t*+1_ + *γ*^2^*r*_*t*+2_ + ⋯ | *s*_*t*_] (i.e. the expected sum of γ-discounted future rewards *r*_*t*_ starting in some state *s*_*t*_). Both in neuroscience and in AI^54^, a typical starting assumption for high-dimensional or continuous tasks is to assume the learner approximates value linearly in some feature basis. That is, it represents the state *s*_*t*_ by a vector of features 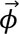 (*s*_*t*_) (henceforth, 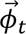) and approximates value as a weighted sum of those features, i.e. *S*_*t*_ = *S*(*s*_*t*_) ≈ 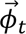 ⋅ 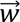. This reduces the problem of value learning (for some feature set) to learning the error-minimizing weights 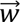 and, more importantly for us, formalizes the state representation itself as a vector of time-varying features 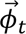. (The linearity assumption is less restrictive than it seems, since the features may be arbitrarily complex and nonlinear in their inputs. For instance, this scheme is standard in AI, including at the final layer in deep-RL models, which first derive features from a video input by multilayer convolutional networks, then finally estimate value linearly from them^55,56^.)

In a standard temporal-difference (TD) learning model, weights are learned by a delta rule using the RPE *δ*_*t*_ = *r*_*t*_ + *γV*(*s*_*t*+1_) − *V*(*s*_*t*_)^57^. A typical cartoon of how these are mapped onto brain circuitry is shown in **Fig. 1a**, with a cortical input population vector for state features projected to scalar value and RPE stages, corresponding, respectively, (since both variables are scalar) to presumed uniform populations of striatal and midbrain DA neurons^1–3^. The RPE then drives weight learning at the corticostriatal synapses via ascending DA projections. Previously proposed models to account for DA heterogeneity often replace the target *r*_*t*_ with one or more additional (potentially non-reward) target outcomes, replicating this scalar prediction model across parallel circuits (“outcome-specific PE models”; **Fig. 1b**)^31–39,58^. These models can explain some types of variation in DA across projection-defined populations.

In contrast, we propose a new “feature-specific RPE model”, which produces heterogeneity even within a projection-defined DA population. This model relaxes the unrealistic anatomical assumption that the corticostriatal stage of this circuit involves complete, uniform convergence from vector state to scalar value (**Fig. 1c**). In fact, projections are substantially topographic at each level of the corticostriatal circuit^43,44,59^. Thus, if different striatal units *i* preferentially receive input from particular cortical features *ϕ*_*i*,*t*_, then value will itself be represented by a distributed feature code *S*_*i*,*t*_ = *ϕ*_*i*_*ϕ*_*i*,*t*_ (see also^2,60^), and, in turn, DA neurons preferentially driven by each “channel” will compute feature-specific reward prediction errors

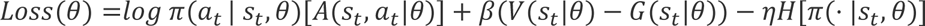

(where *N* is a scale factor equaling the number of channels). Importantly, due to linearity, the aggregate response (summed over channels *i*) at the value stage reflects the original scalar value, and the aggregate response at the RPE stage corresponds to the original scalar RPE.

Thus (assuming the ascending dopaminergic projection is sufficiently diffuse as to mix the channels prior to the weight update), the model corresponds algebraically to the classic one, just mapped onto the brain circuitry in a more realistic way. Our primary insight is that nonuniform convergence from the state layer to the value layer, and from the value layer to the dopamine layer, will result in value and RPE stages that reflect input feature variation, but preserve the scalar RPE model (**Fig. 1a**) due to linearity. Note that this insight holds, at least approximately, under more realistic models in which channels partially mix at each step due to anatomically limited convergence (so that, for instance, a feature at the value or RPE stage is the average of several features from the afferent population) or in which linearity is only approximate.

While still quite stylized, this account represents a more realistic picture compared to the traditional scalar story of both value and RPE. Notably, it better reflects the known topography in the projections from cortical inputs to MSNs and from MSNs to dopamine units^44,45,49^, and physiological findings that both input populations show variable tuning to similar features as do the DA neurons^5,61–66^. Incorporated in the feature-specific RPE model (even without specifying a particular feature basis), these properties directly imply several aspects of the heterogeneous dopamine response. Consider times when primary reward is not present, such as during the navigation and decision-making period in Engelhard et al.^4^ Here, *r*_*t*_ = 0 and Equation 1 reduces to *δ*_*i*,*t*_ = *ϕ*_*i*_(*γϕ*_*i*,*t*+1_ − *ϕ*_*i*,*t*_): that is, each DA unit reports the time-differenced activity in its feature, weighted by its own association *ϕ*_*i*_ with value. Depending on what the features 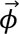 are, this would explain dopaminergic response correlations with different, arbitrary covariates: to the extent some feature *ϕ*_*i*_ is task-relevant, it will have nonzero *ϕ*_*i*_ (where the sign of *ϕ*_*i*_ is determined by *ϕ*_*i*_’s partial correlation with value in the presence of the other features), and its derivative will correlate with a subset of neurons. Even objectively task irrelevant features *ϕ*_*i*_ might transiently or incidentally correlate with value, producing responses to different such features in different subsets of neurons.

Conversely, at outcome time, all modeled RPE units are likely to respond differentially for reward than nonreward (due to *r*_*t*_ being shared across all channels in Equation 1) as in Engelhard et al^4^. This reward response may also be modulated by its predictability, due to each channel’s share of the temporal difference component, *ϕ*_*i*_(*γϕ*_*i*,*t*+1_ − *ϕ*_*i*,*t*_), but is unlikely to be completely predicted away by most individual features. Finally, since the standard RPE *δ*_*t*_ is equal to the sum over all channels *Σ*_*i*_*δ*_*i*,*t*_ by construction, the model explains why neuron-averaged data (or bulk signals as in fiber photometry or BOLD), as often reported, resemble TD model predictions even potentially in the presence of much inter-neuron variation.

### Deep RL network to simulate the Feature-Specific PE model

Although our feature-specific PE model is fully general, in order to simulate the model in the context of a specific task, we need to specify an appropriate set of basis functions for the task state. We consider our previously reported experiment in which mice performed an evidence accumulation task in a virtual reality environment while VTA DA neurons were imaged^4^ (**Fig. 2a**). In this task, mice navigated in a virtual T-maze while viewing towers that appeared transiently to the left and right, and were rewarded for turning to the side where there had been more towers.

**Figure 2:**
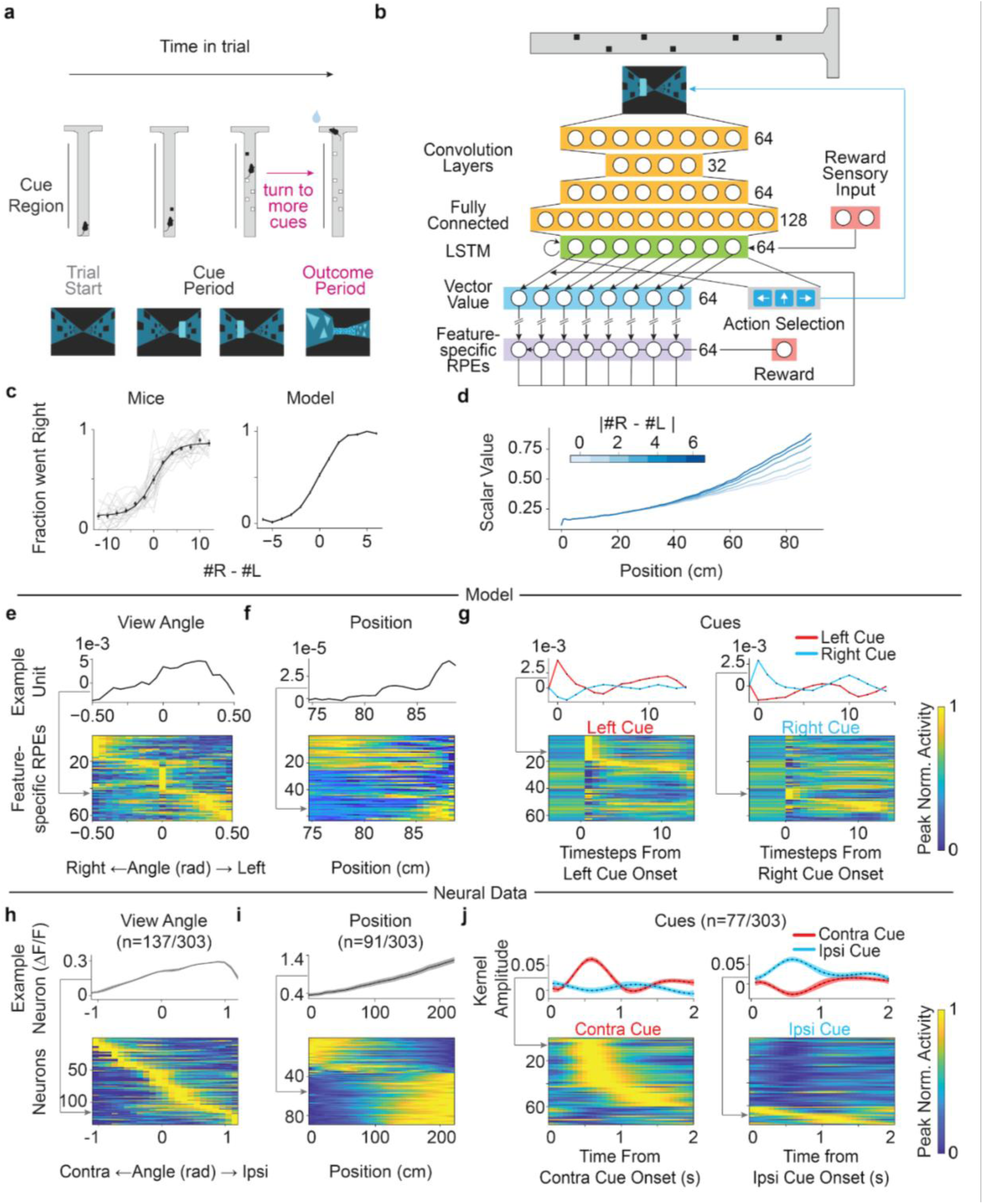
Feature-specific RPEs derived from a deep reinforcement learning model trained on a VR evidence accumulation task are heterogeneously modulated by behavioral variables during the navigation and decision period, similar to DA neurons. **(a)** Task schematic of the VR task in which mice accumulated visual evidence (cues) as they ran down the stem of a T-maze and were rewarded if they turned to the side with more cues at the end. Video frames of the maze are shown below each maze schematic. **(b)** A deep reinforcement learning (RL) network took in video frames from the VR task, processed them with 3 convolution layers, a fully connected layer (orange), and an LSTM layer (green), and outputted an action policy (gray), which inputted the chosen action back into the VR system (blue arrow). The second to last layer of the deep RL network (the LSTM layer) and the weights for the critic served as the inputs to form the feature-specific RPEs (purple). Reward information (red) was inputted both as a one-hot sensory signal to the LSTM layers (for rewarded and unrewarded trials) and also to the feature-specific RPEs. **(c)** Psychometric curves showing the mice’s performance (left) and agent’s performance (right) after training. The fraction of right choices is plotted as a function of the difference of the right and left towers presented on the trial. For the mice, gray lines denote the average psychometric curves for individual sessions and the black line denotes a logistic fit to the grand mean with bars denoting the s.e.m. (N = 23 sessions). For the model, black bars indicate the s.e.m. (N = 5000 trials) **(d)** Deep RL model’s scalar value (sum over units in vector value) during the cue period decreased as trial difficulty (measured by absolute value tower difference, blue gradient) increased. **(e)** Feature-specific RPE unit activity with respect to the view angle of the agent (min-max normalization; top: example unit; bottom: all units). **(f)** Same as **(e)** but for position of the agent in the final 15 cm of the maze. **(g)** Same as (**e-f**) but for left (red) and right (blue) cues. **(h-j)** Same as **(e-g)** but for the subset of neurons from Engelhard et al^4^ tuned to **(h)** view angle of the mice, **(i)** position, and **(j)** contralateral (red) and ipsilateral (blue) cues. Fringes represent ±1 s.e.m. In **(h-i)** peak normalized ΔF/F signals are plotted while in **(j)** cue kernels are plotted from an encoding model used to quantify the relationship between the behavioral variables and each neuron.

To simulate the vector of feature-specific PEs (in this task, RPEs), we took advantage of the fact that this task was based in virtual reality, and therefore we could train a deep RL agent on the same task that the mice performed to derive a vector of features, and in turn, a corresponding distributed code for values and RPEs (**Fig. 2b**). In particular, we used a deep neural network to map the visual images from the virtual reality task to 64 feature units (via three convolutional layers for vision, then a layer of LSTM recurrent units for evidence accumulation; see Methods). These features were then used as common input for an actor-critic RL agent: a linear value-predicting critic (as above, producing the feature-specific RPE that sums to the traditional scalar RPE) and a softmax advantage learner responsible for choosing an action (left, right, forward) at each step. We trained the network to perform the task using the A2C algorithm^56^. Note that although A2C does not use the feature-specific RPEs individually for training, only their sum, this is also true in our model: the decomposition over features, in both cases, arises at an intermediate stage of computation.

After training, the agent accumulated evidence along the central stem of the maze to ultimately choose the correct side with accuracy similar to mice (shown as a psychometric curve in **Fig. 2c; Extended Data Fig. 1** shows that this and later results are consistent in an independent training run). A minimal abstract state space underlying the task is 2D, consisting of the position along the maze and the number of towers seen (on left minus right) so far. When examining average responses of the trained state features from the network in this space, we find that units are tuned to different combinations of these features, implying that they collectively span a relevant state representation for the task (**Extended Data Fig. 2**). Further, the scalar value function output by the trained agent while traversing the maze, derived by summing the value vector, is modulated by trial difficulty (operationalized, following the mouse study, as the absolute value of the difference in the number of cues presented on either side), meaning that the trained agent can predict the likelihood of reward on each trial (**Fig. 2d**).

### Feature-specific RPEs have heterogeneous selectivity during the cue period

The key finding from our prior cellular resolution imaging of DA neurons during the virtual T-maze task was heterogenous coding of task and behavioral variables during the navigation and decision period, such as view angle, position, and cue side (contralateral versus ipsilateral)^4^. This heterogeneity was followed by relatively homogeneous responses to reward during the outcome period.

We first sought to determine if the feature-specific RPEs from our agent had heterogenous tuning to various behavioral and task variables during the cue period, similar to our neural data. We first considered the view angle during the central stem of the maze, which had no effect on reward delivery in the simulated task, but which the agent could nonetheless rotate by choosing left or right actions while in the stem of the maze. The feature-specific RPEs displayed idiosyncratic selectivity across units for the range of possible view angles (**Fig. 2e**), which qualitatively resembled our previous results from DA neuron recordings (**Fig. 2h**). A subset of RPE units showed position selectivity, including both downward and upward ramps towards the end of the maze (**Fig. 2f**), again qualitatively resembling our neural recordings (**Fig. 2i**). Finally, as in the neural data (**Fig. 2j**), our feature-specific units also had idiosyncratic and heterogeneous cue selectivity, including preference for right vs left cues and diversity in the timing of the response (**Fig. 2g**). Similar to the neural data^4^, we observe clustering of these feature responses (**Extended Data Fig. 3a**), with different clusters responding most strongly to cues, position, and choice, respectively (**Extended Data Fig. 3b**).

### Reward-irrelevant responses in the feature-specific RPEs

A property of the neural data is the encoding of task features that appear to be reward-irrelevant^4^. In the model, the requirement of the network to extract relevant task state from the high-dimensional video input implies the possibility that reward-irrelevant aspects of the input may “leak” into the state features, and ultimately into the feature-specific RPEs – even if they average out in the scalar RPE. Although we of course do not intend backpropagation as a mechanistic account of how the brain learns features, its use here exemplifies the more general principle that a low-dimensional output objective (choosing actions and predicting scalar reward) imposes few constraints on higher-dimensional upstream feature representations.

To test the validity of this idea, we sought to examine whether there may be coding of reward-irrelevant visual information in the feature-specific RPEs of the agent. Because the reward-irrelevance of many task variables is difficult to definitively specify, we focused on unambiguously reward-irrelevant visual structure in the task, namely incidental background patterns that appear on the wall in the stem of the maze (**Fig. 3a**). This background pattern repeats every 43 cm, a structure which is clearly visible as off-diagonal banding in the matrix of similarity between pairs of video frames across all combinations of locations. The structure is also visible as peaks in the 1D function (similar to an autocorrelation) showing the average similarity as a function of the distance between frames (**Fig. 3b**). To investigate whether the same irrelevant feature dimensions are present at the level of the feature-specific RPEs, we repeat the same analysis on them. (To ensure the network inputs actually reflect such repeating similarity structure, for this analysis we exposed the trained network to a maze traversal with a fixed view angle). The resulting feature-specific RPEs show the same pattern of enhanced similarity at the characteristic 43 cm lag, supporting the expectation that task-irrelevant features do propagate through the network (**Fig. 3c-d**). This particular effect remains a prediction for future neural experiments: because the spatial frequency of the irrelevant background patterns was so high, the patterns cycled too quickly (more than once per second at typical running speed) for any correlations with DA responses to be resolved given the temporal properties of the measurements (which is an issue in empirical but not simulated data. However, it suggests a straightforward design for future experiments, using slower irrelevant stimuli. Note that this pattern is not present in the scalar RPE, indicating that the loss function during training eliminates task-irrelevant features from the aggregated but not the individual RPEs (**Extended Data Fig. 4**).

**Figure 3:**
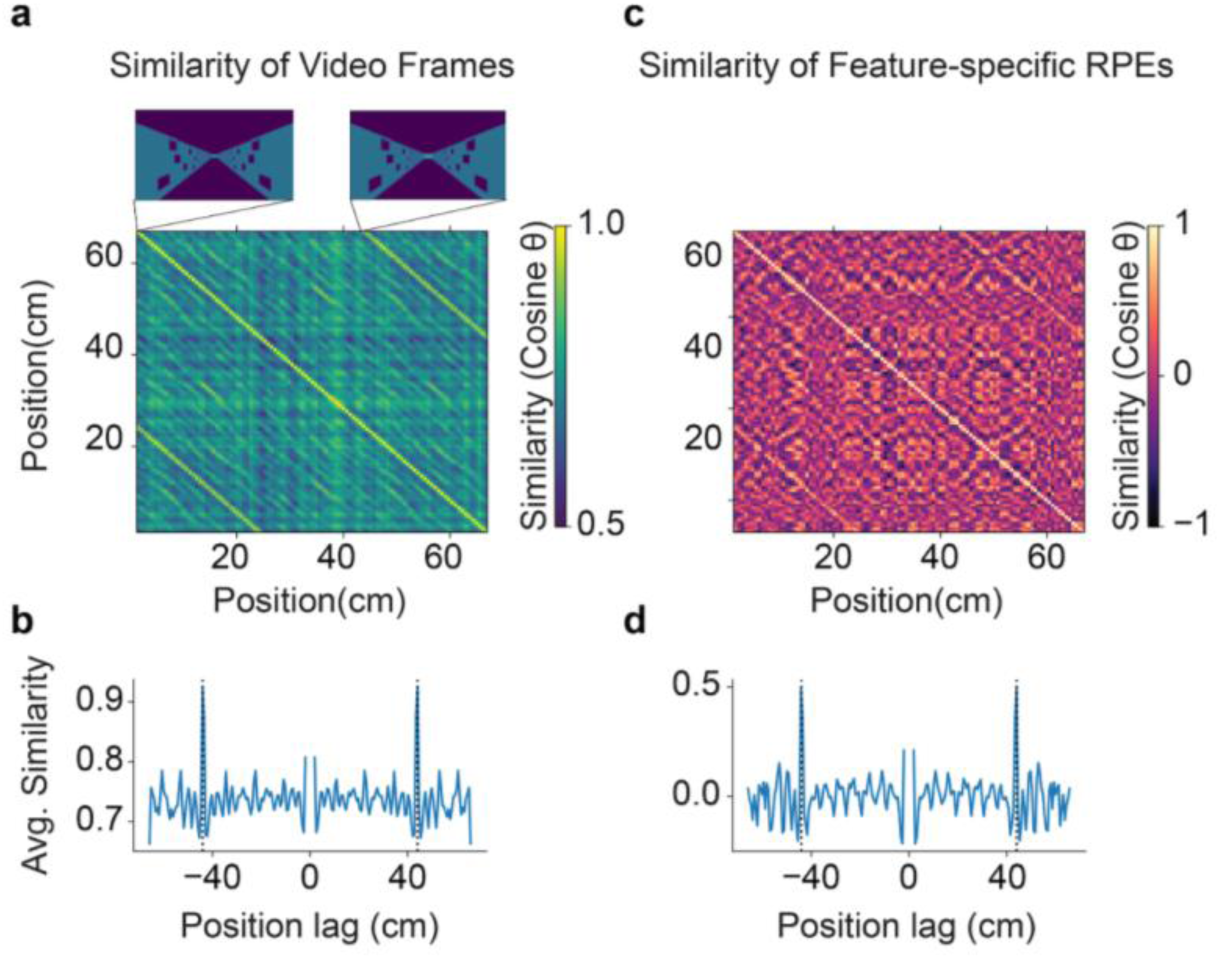
Feature-specific RPEs reflected incidental high-dimensional visual inputs. **(a)** Similarity matrices of the video frames, which measure the similarity between pairs of video frames (quantified by the cosine of the angle between them when flattened to vectors) across different position combinations. The off-diagonal bands correspond to the wall-pattern repetitions (see video frames for position 0cm and 43cm at insets above). **(b)** Average similarity as a function of distance between frames, indicating that the average similarity peaked at the same position lag (43 cm) for video frames. **(c-d)** same as **(a-b)**, but with feature-specific RPEs.

### Cue responses are consistent with feature-specific RPEs

While our feature-specific RPE model implies idiosyncratic and even task-irrelevant tuning in individual DA neurons, it also makes a fundamental prediction about the nature of these responses. In particular, responses to individual features represent not generic sensory or motor responses but feature-specific components of RPE. What in practice this means for a given unit depends both on what is its input feature *ϕ*_*i*_, and also what other task features the brain represents (since net reward prediction and thus RPE are jointly determined by multiple cues). In general, we would expect that units that appear to respond to some feature (such as contralateral cues) do not simply reflect sensory responses (the presence of the cue) but rather should be further modulated by the component of RPE elicited by the feature. This could be particularly evident when considering the response averaged over units selective for a feature.

To test this hypothesis, we performed a new analysis, by subdividing cue-related responses (which are largely side-selective in both the model and neural data, **Fig. 2g, j**) to determine if they were, in fact, additionally sensitive to the prediction error associated with a cue on the preferred side. For this, we distinguished these cues as *confirmatory* – those that appear when their side has already had more cues than the other and therefore (due to the monotonic psychometric curve, **Fig. 2c**) are associated with an increase in the probability that the final choice will be correct and rewarded, i.e. positive RPE – vs *disconfirmatory* – cues whose side has had fewer towers so far and therefore imply a decreased probability of reward (**Fig. 4a**). As expected, when we considered the population of cue-onset responding feature-specific RPE units from the deep RL agent, these responses were stronger, on average, for confirmatory than disconfirmatory cues (**Fig. 4b**), reflecting the component of RPE associated with the cue. We next reanalyzed our previous DA recordings based on this same insight. Consistent with the hypothesis that these cue responses reflect partial RPEs for those cues, the responses of cue-selective DA neurons were much stronger for confirmatory than disconfirmatory cues (**Fig. 4c**). This implies that the heterogenous cue selectivity in DA neurons is indeed consistent with a cue-specific RPE. Importantly, the fact that these cue-responsive neurons are overwhelmingly selective for contralateral cues implies that these responses, combined across hemispheres, simultaneously represent two separate components of a vector of multidimensional feature-specific RPEs (as opposed to a classic scalar RPE). We confirm this by further separating the neural response between neurons recorded from each hemisphere (**Extended Data Fig. 5a,b**). As in the model (**Extended Data Fig. 5c,d**), responses to cues contralateral to each side both distinguish confirmatory from disconfirmatory cues.

**Figure 4:**
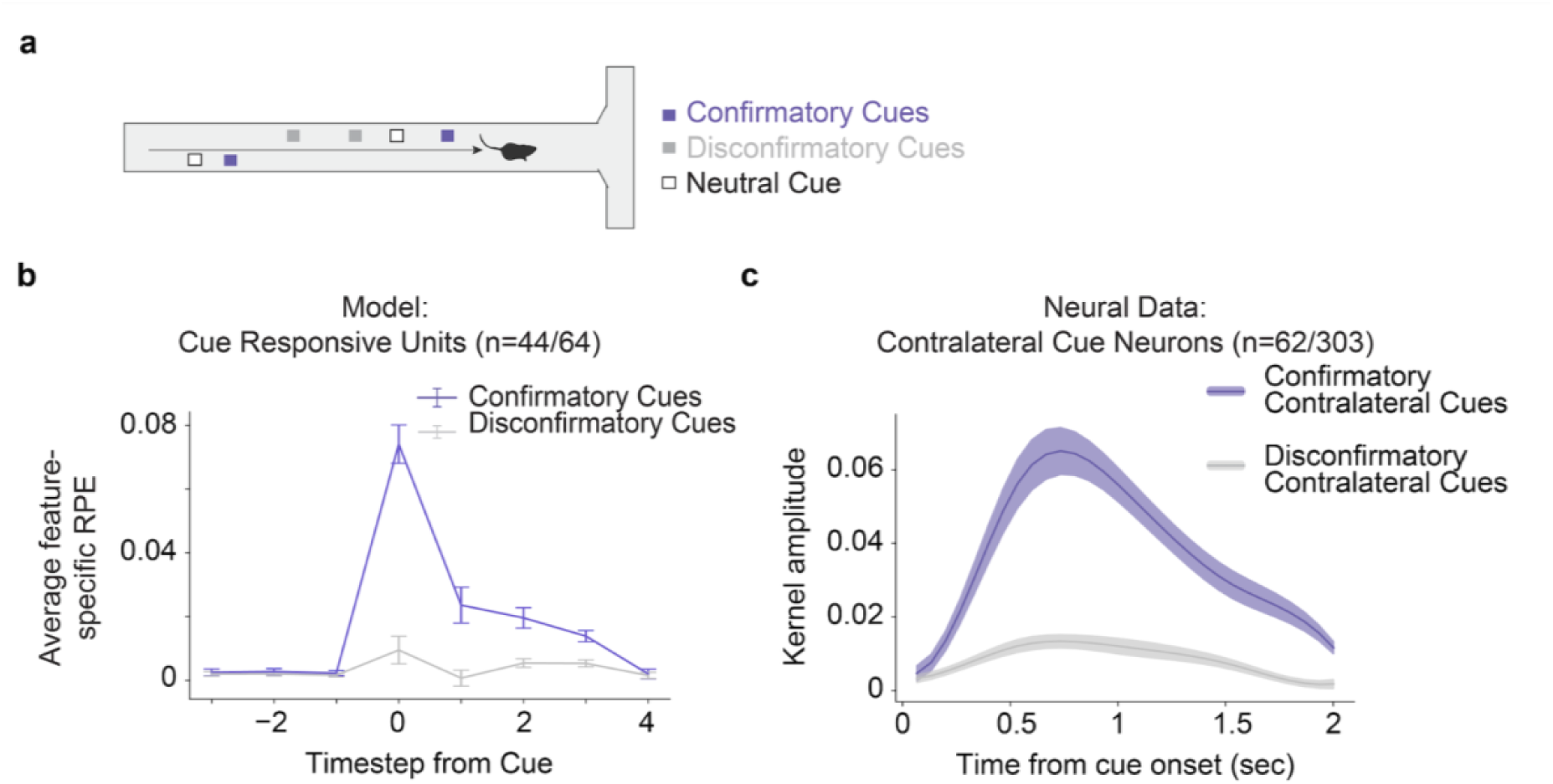
Cue responses in model and DAergic neurons reflected RPEs with respect to cues, rather than simply their presence. **(a)** Example trial illustrating confirmatory cues (purple), defined as cues that appear on the side with more evidence shown thus far, and disconfirmatory cues (gray), which are cues appearing on the side with less evidence shown thus far. Neutral cues (white) occur when there has been the same amount of evidence shown on both sides. **(b)** Average response of feature-specific RPE units modulated by cues (N = 44/64) to confirmatory (purple) and disconfirmatory cues (gray). Error bars indicate ±1 s.e.m. **(c)** Average responses of the contralateral cue responsive DA neurons for confirmatory (purple) and disconfirmatory (gray) contralateral cues. Colored fringes represent ±1 s.e.m. for kernel amplitudes (N = 62 neurons, subset of cue responsive neurons from Fig. 2j that were modulated by contralateral cues only).

### Uniform responses to reward at outcome period

In addition to explaining heterogeneity during the navigation and decision-making period, the feature-specific RPE model also explained the homogeneity of the neural responses to reward during the outcome period. Reflecting the standard properties of an RPE, the simulated scalar RPE (averaged over units) responded more for rewarded than unrewarded trials (**Fig. 5a**). Since this aspect of the response ultimately arises (in Equation 1) from a scalar reward input, it is highly consistent across the units (**Fig. 5b**), matching the neural data from our experiment (**Fig. 5c**) and the widely reported reward sensitivity of DAergic units.

**Figure 5:**
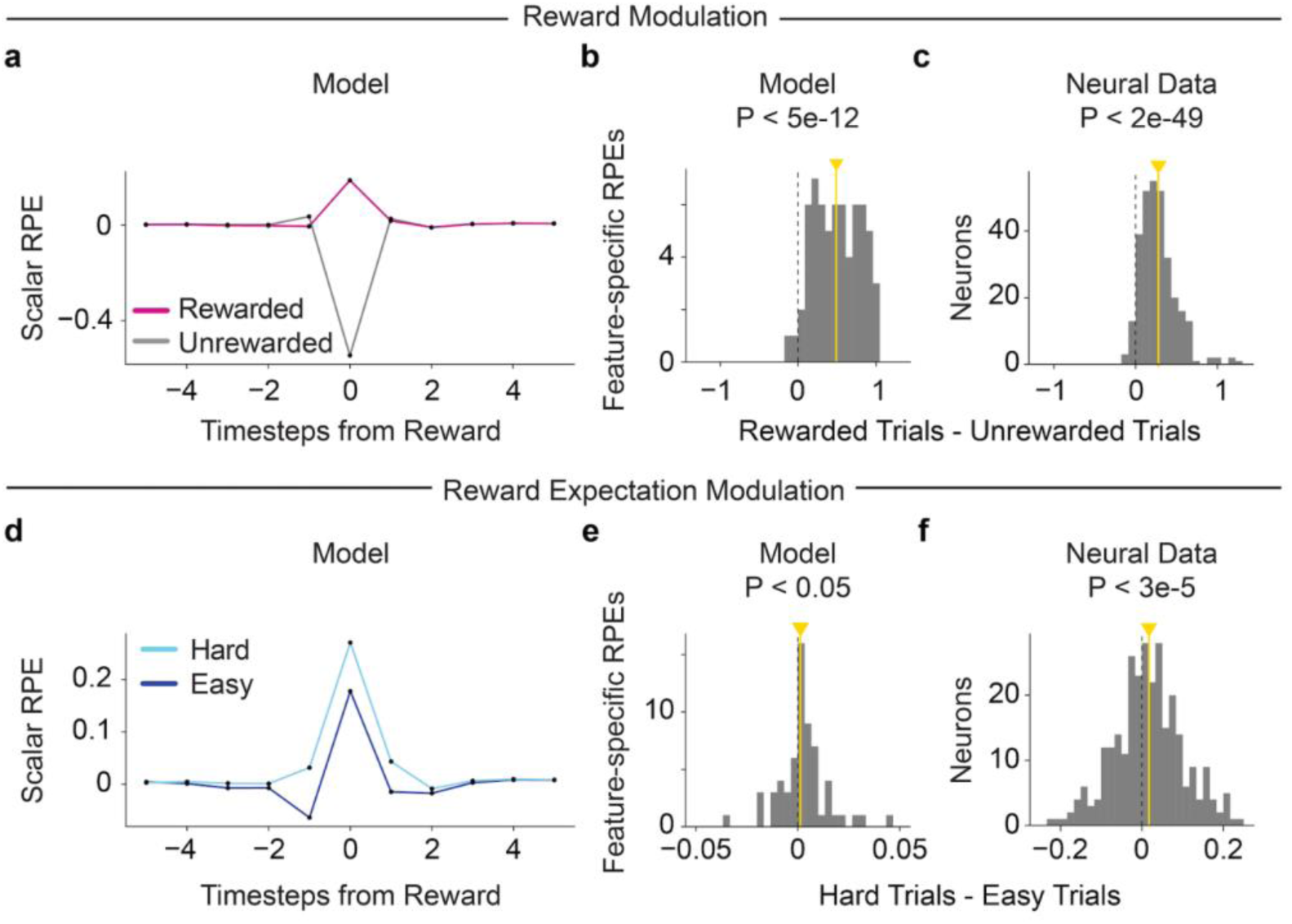
Feature-specific RPE units and DAergic neurons consistently respond to reward, but show heterogeneous modulation by reward expectation. **(a)** Scalar RPE (sum of feature-specific RPEs units) time-locked at reward time for rewarded (magenta) minus unrewarded trials (gray). **(b)** Histogram of feature-specific RPE units’ response to reward minus omission at reward time (P < 5e-12 for two sided Wilcoxon signed rank test, N = 64). Yellow line indicates median. **(c)** Same as **(b)**, but with all imaged DA neurons^4^, using averaged activity for the first 2 seconds after reward delivery, baseline corrected by subtracting the average activity 1 second before reward delivery (P < 2e-49 for two sided Wilcoxon signed rank test, N = 303). **(d)** Scalar RPE time-locked at reward time for rewarded trials, split by hard (light blue) and easy (dark blue) trials, defined as trials in the bottom or top tercile of trial difficulty, measured by the absolute value of the difference between towers presented on either side. **(e)** Histogram of feature-specific RPE units’ responses to difficult minus easy rewarded trials at reward time (P < 0.05 for two sided Wilcoxon signed rank test, N = 64) with median indicated by yellow line. There is an outlier data point at 0.29 for a feature-specific RPE unit showing strong reward expectation modulation. **(f)** Same as **(e)** but with the averaged activity of DA neurons^4^ corrected by their baseline activity 1 second before reward delivery (P < 3e-5 for two sided Wilcoxon signed rank test, N = 303).

Equation 1 also implies a subtler prediction about the feature-specific RPEs, which is that although the modulation by reward is largely uniform across units, the modulation of this outcome response by the reward’s predictability should be much more variable. This is because this latter modulation arises from the value terms in Equation 1, which are distributed across features (i.e., value is represented as a vector over features). In this task, reward expectation can be operationalized based on the absolute difference in tower counts, which is a measure of trial difficulty (predicting the actual chance of success as shown in **Fig. 2c**), That is, when a reward occurs following a more difficult discrimination, the agent will have expected that reward with lower probability (**Fig. 2d**). Accordingly, the simulated scalar RPE was larger for rewards on hard than easy trials (**Fig. 5d**; **Extended Data Fig. 6**). However, when broken down unit-by-unit, although the median of individual units in the feature-specific RPE model was consistent with the scalar RPE (P < 0.05 two-sided Wilcoxon signed rank test for N=64), this effect varied widely across units (**Fig. 5e**). A similar finding emerged from the neural data: while on average the reward response across reward-responsive population was modulated as expected by expectation, there was high variability in the modulation across units (**Fig. 5f**).

### The successor representation model cannot account for the reward-predictive aspects of VTA DA heterogeneity

Another candidate account of VTA DA heterogeneity is the successor representation (SR) model (**Fig. 6a**)^39^, which is an example of an outcome-specific PE model (**Fig. 1b**). It posits that DA heterogeneity can arise from the prediction of different sensory features in different parallel circuits, rather than (or in addition to) prediction of reward. We simulated a set of SR error signals, by training a successor representation over the features produced by the DeepRL network trained on the task and then examining sensory prediction errors over these predictions. Similar to the neural data and to the feature-specific RPE model, the SR model produced heterogeneous responses during cue period (**Fig. 6b-d**). However, these modulations were purely sensory rather than value-driven, and thus did not replicate the neural data. For example, responses to disconfirmatory cues were stronger than confirmatory cues (**Fig. 6e**, opposite to the data and the feature-specific RPE model), because disconfirmatory cues are on average more surprising given the task statistics. Similarly, and contrary to the hallmark feature of VTA DA neurons, the SR model’s outcome-period responses are not consistently modulated by reward, showing no difference for averaged sensory PE during rewarded and unrewarded trials (**Fig. 6f, h**) and also no consistent modulation by reward expectation (**Fig. 6g, i**).

**Figure 6:**
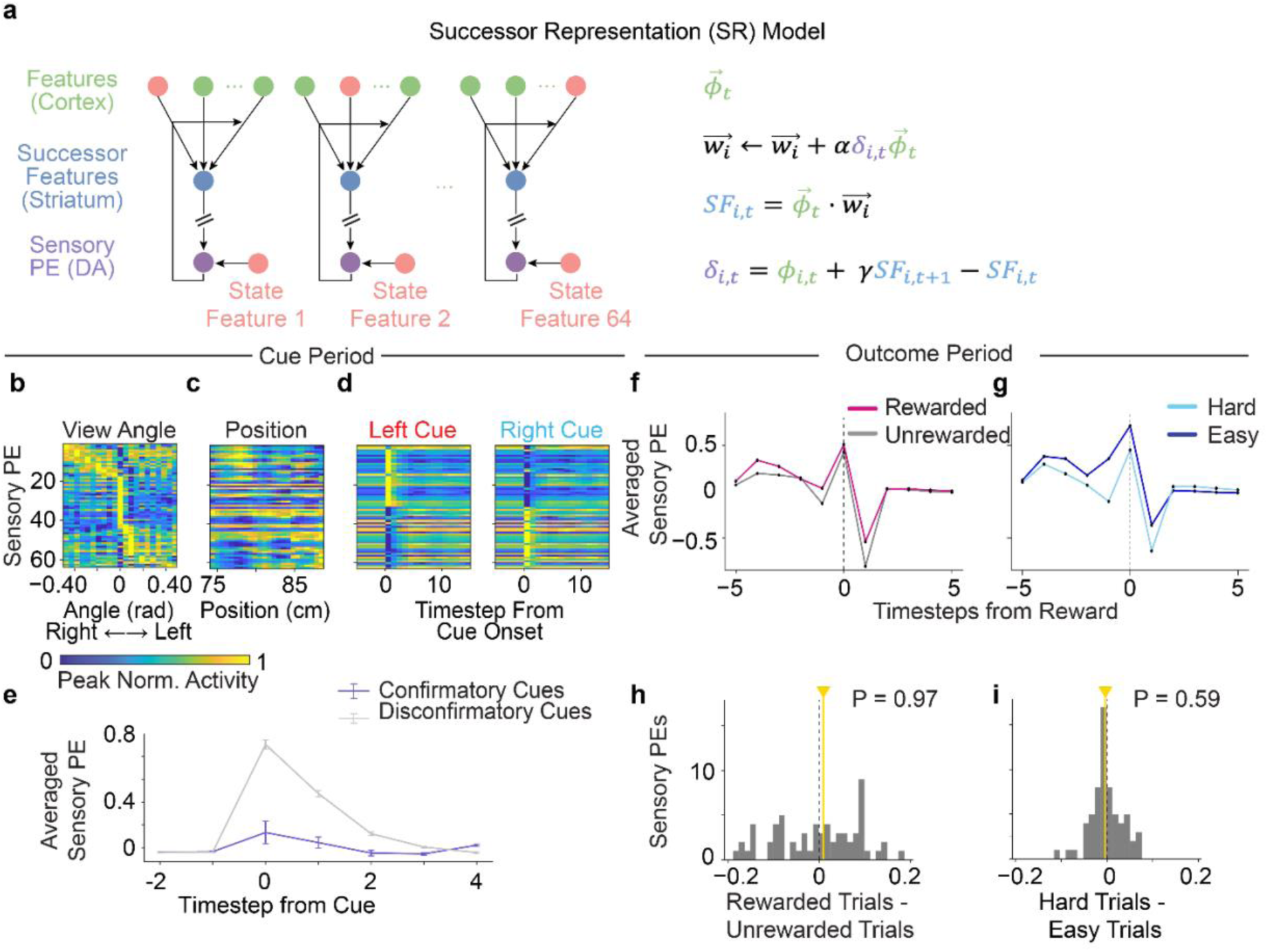
Sensory prediction errors from the successor representation model are heterogeneous during the cue period, but are not responsive to confirmatory cues and reward. **(a)** Schematic of the successor representation model to produce sensory prediction errors. The model was trained on the same state features as the feature-specific RPE model (green), with a vector of successor features (blue) and vector of sensory PEs (purple) with respect to each of the 64 features. **(b)** Sensory PEs plotted with respect to view angle. Each row corresponds to a unit’s peak-normalized response to view angle averaged across trials. **(c-d)** Same as **(b)** but for **(c)** position and **(d)** left (red) and right (blue) cues. **(e)** Sensory PE averaged across units in response to confirmatory (purple) and disconfirmatory cues (gray) shows a stronger response for disconfirmatory cues. Error bars indicate ±1 s.e.m. **(f)** Sensory PE averaged across units split by rewarded (magenta) and unrewarded (gray) trials (at the time of reward. **(g)** Same as **(f)** but for easy trials (dark blue) and hard trials (light blue) instead, defined as in Fig. 5d. **(h)** Histogram of the difference in sensory PE units’ responses at reward time for rewarded minus unrewarded trials, with the yellow line indicating the median (P = 0.97 for two sided Wilcoxon signed rank test, N = 64). **(i)** Same as **(h)** but for easy versus hard trials. Only rewarded trials are included (P = 0.59 for two sided Wilcoxon signed rank test N = 64).

### Distributional RL cannot account for VTA DA heterogeneity during the cue period

Another prominent example of an outcome-specific PE model is the distributional RL model (**Fig. 7a**)^33^, which posits that parallel channels predict different expectiles of the distribution of expected future reward, i.e. different weightings of better vs. worse outcomes. This model has been shown to predict subtle variability in cue and outcome DA responses in classical conditioning^33^, but it is unclear to what extent it might also capture the broader range of variability in more complex tasks like the one studied here. We simulated expectile PE signals based on the features produced by the DeepRL network trained on the task. Since distributional RL uses a common reward outcome across all channels, it was able to capture many of the reward-related aspects of the neural data, including stronger responses to confirmatory over disconfirmatory cues (**Fig. 7e**) and uniform response to reward (**Fig. 7f,h**). It was also able to produce some heterogeneity over responses during the cue period (**Fig. 7b-d**): units did vary in their response to features like cues or view angle, ultimately because these are associated with subtly different outcome distributions. However, because it combines all features for each prediction error, the model predicts symmetric responses to features that are equivalent with respect to outcomes: e.g., cues on a particular side, or view angles on the right versus left. The prominent side-selectivity of DA responses thus suggests there is additional variability across neurons related to their specific feature inputs, over and above that accounted for by their associated outcome expectiles.

**Figure 7:**
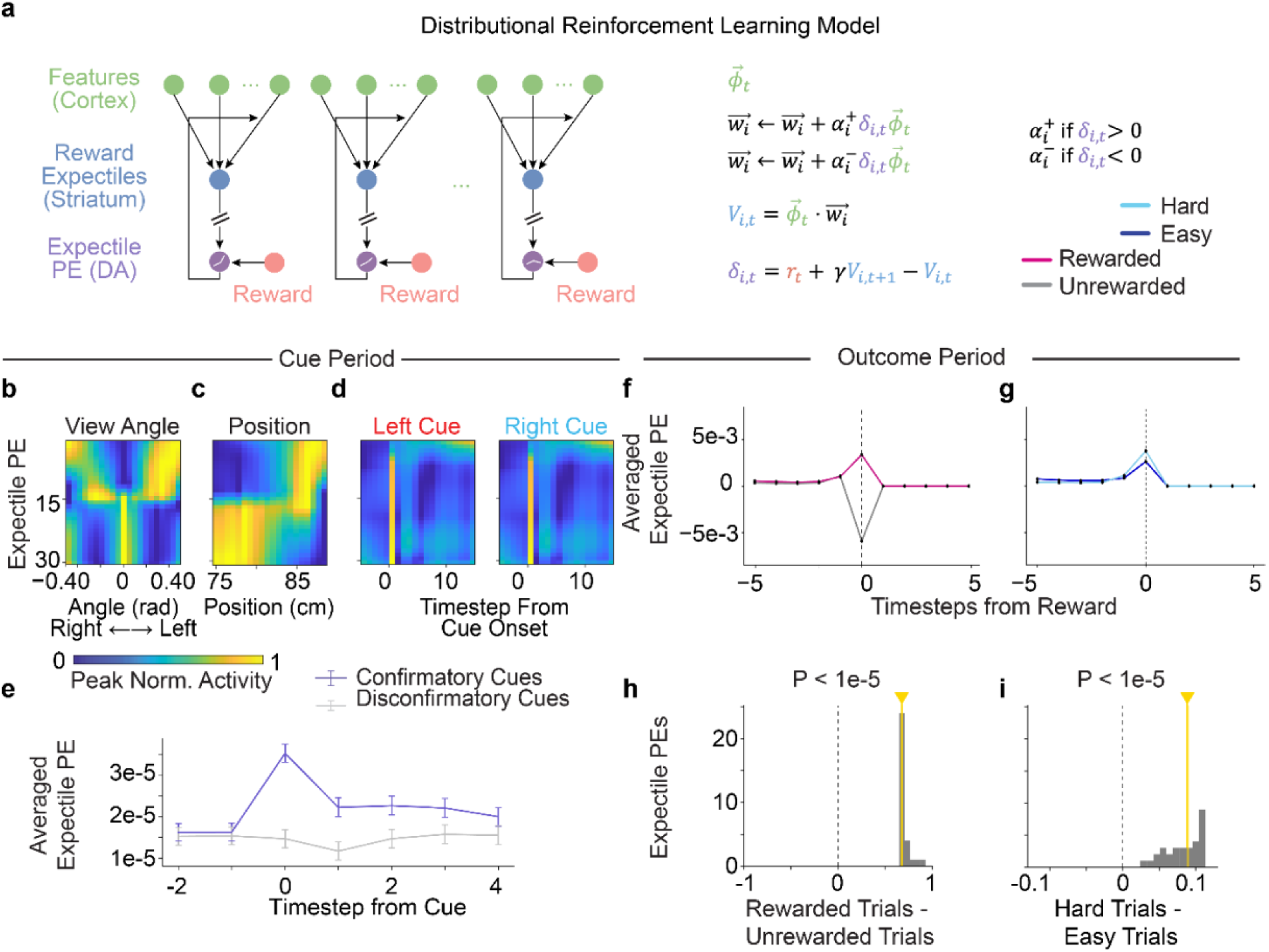
Expectile prediction errors from a distributional RL model are not heterogeneous during the cue period. **(a)** Schematic for the distributional reinforcement learning model which produces expectile prediction errors. The model uses the same state features as the feature-specific RPE model (green), with all the features input into 31 channels. These channels represent different expectiles of value (evenly spaced between 0 and 1), and 31 corresponding expectile prediction errors, which differentially weight positive and negative information. **(b)** Expectile PE activity plotted with respect to view angle, peak normalized for each unit and sorted in the original distributional channel order. **(c-d)** Same as **(b)** but with **(c)** position and **(d)** left (red) versus right (blue) cues. **(e)** Expectile PEs averaged across units, responding to confirmatory (purple) and disconfirmatory (gray) cues. Error bars indicate ±1 s.e.m. **(f)** Expectile PEs averaged across units time-locked to reward, split by rewarded (magenta) versus unrewarded (gray) trials. **(g)** Same as **(f)** but split by easy trials (dark blue) and hard trials (light blue). Only rewarded trials are included, and trial difficulty is defined the same as Fig. 5d. **(h)** Histogram of expectile PEs at reward time, showing the difference in their response for rewarded and unrewarded trials (P < 1e-5 for two-sided Wilcoxon signed rank test, N = 31 units). Median response shown with the yellow line. **(i)** Same as **(h)**, but for trial difficulty, taking the difference between responses for hard and easy rewarded trials (P < 1e-5 for two-sided Wilcoxon signed rank test, N = 31 units).

### Feature-specific action prediction errors explain DA heterogeneity in SNc

So far we have shown that the feature-specific PE model captures, and outcome-specific PE models generally fail to match, heterogeneity of DA responses within VTA. This is consistent with our expectation that outcome-specific PEs are likely more relevant to heterogeneity between rather than within a DA subpopulation. To what extent does each class of model explain data from other tasks and other DA populations? Recent work has shown that DA neurons in the substantia nigra pars compacta^7,13,21,67^ (SNc) and the substantia nigra pars lateralis^40^ (SNl) can exhibit qualitatively different response properties than those in the VTA, and that these differences may relate to their striatal projection target. In particular, some SNc/SNl DA neurons are tuned to actions, for example displaying a transient increase in firing shortly before action onset followed by a longer depression throughout action execution^5,21,67–69^. Interestingly, as a general rule, action selectivity in DA responses in SNc tends also to be accompanied by reduced response to reward^7,21,40,67^. For example, Parker et al.^7^ recorded DA terminals in dorsomedial striatum (likely arising from SNc) during a reversal learning lever-choice task (**Fig. 8a**) and showed that they responded robustly to contralateral movements (e.g., press of the lever contralateral to the recording site), but (in contrast to classic RPEs observed in nucleus accumbens, NAc, a VTA target) showed little modulation to reward consumption or reward-predicting cues.

**Figure 8:**
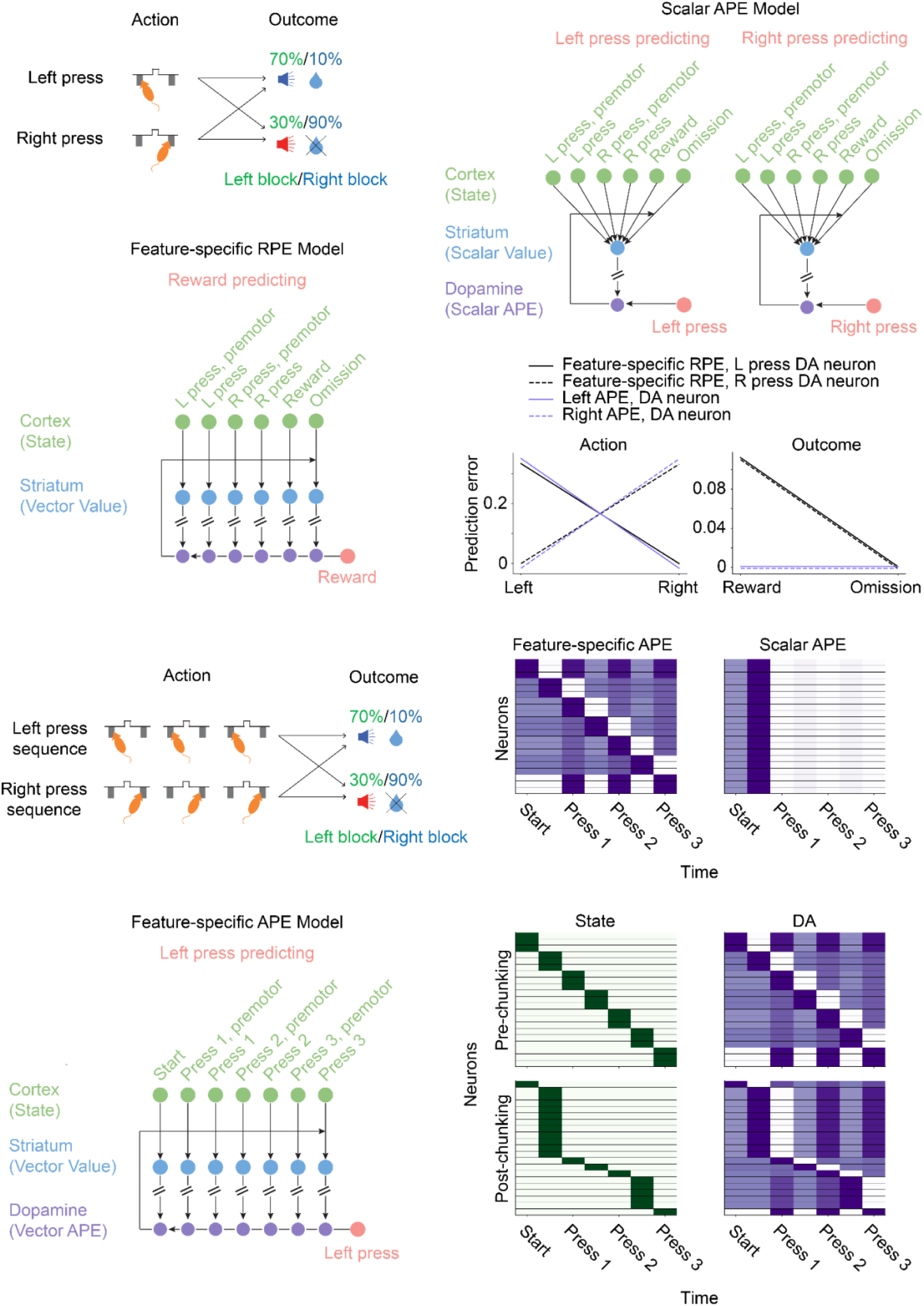
Feature-specific action prediction errors provides a potential explanation for movement-related heterogeneity in SNc DA neurons. **(a)** Schematic for the Parker et al^7^ reversal task. Mice were offered a choice between two levers, each of which provided reward with some probability. During a left block (green), the left lever rewarded with probability 0.7 and the right lever rewarded with probability 0.1. During a right block (blue), this contingency was reversed. Identity of the current block (left or right) was unsignaled to the mice and swapped with a pseudorandom schedule. **(b)** Schematic of the scalar APE model. The model was trained with features encoding a one-hot representation of the state (green), scalar action value (blue), and scalar APE (purple). The left schematic displays the left-preferring subcircuit and the right schematic displays the right-preferring subcircuit. **(c)** Schematic of the feature-specific RPE model used on the Parker et al reversal learning task^7^. Input features were the same as the scalar APE model in **(b)**. The value layer (blue) and the RPE layer (purple) were feature-specific. **(d)** Dopaminergic predictions from the scalar APE model (purple) and the feature-specific RPE model (black) during action execution (left) and outcome (right) in the reversal learning task^7^. **(e)** Schematic for a simplified version of the Jin and Costa task^5^. Mice are offered a choice between two levers, each of which provided a reward with some probability after being pressed three times. Reward contingencies are the same as **(a)**. **(f)** Schematic for the left subcircuit of the feature-specific APE model used on the Jin and Costa^5^ task. The model was trained with features encoding the state (green), vector action value (blue), and feature-specific APEs. **(g)** Activity of the DA neurons in the feature-specific (left) and scalar (right) APE models during the task. Neurons are sorted by peak onset. Black horizontal lines separate the neuronal activity profiles. **(h)** Activity of the state (green) and DA neurons (purple) in the feature-specific APE model before (top) and after (bottom) chunking. Black horizontal lines separate the neuronal activity profiles.

Can outcome-specific PE or feature-specific RPE models explain the apparent tradeoff, across regions, between movement and reward selectivity? We turn our attention to the action prediction error model^40–42^ (APE) as a potential explanation. Briefly, the APE model is a type of outcome-specific prediction error model comprised of parallel circuits that each predict different movements (e.g. left vs. right lever press), instead of reward (**Fig. 8b**, see Methods).

We compared the results of this model and our feature-specific RPE model (**Fig. 8c**) in simulations of the Parker et al.^7^ probabilistic reversal task using a hand-designed “one-hot” state/feature space (**Extended Data Fig. 7a**, see Methods). We examined simulated DA responses from both models during choice and outcome, specifically focusing on the DA neurons from the left and right subcircuits in the APE model and the DA neurons associated with the left and right choice features in the feature-specific RPE model (**Fig. 8d**). At choice, these make roughly similar predictions: the DA neurons fire more when their preferred action is made. At outcome, however, their predictions differ. Due to the *r*_*t*_/*N* term in **Equation 1**, the feature-specific RPE DA neurons emit a fixed-magnitude, positive response to reward, whereas the APE DA neurons do not. As such, while the feature-specific RPE model can produce DA neurons tuned to different movements, it does not explain the tradeoff between action selectivity and reward selectivity. The decoupling of actions from rewards suggests that for understanding why SNc-DMS responses differ from VTA-NAc ones, the principles of outcome-specific PE apply.

However, even within the movement-selective population in the SNc, there is additional functional heterogeneity. For example, Jin and Costa^5^ recorded individual DA neurons during a task in which mice had to sequentially press a lever multiple times for reward, and reported most DA neurons had specific preferred presses (e.g., the first or last presses within a sequence). Differential press-specific tuning is difficult to explain with a classic APE model. Since later presses are predicted by earlier presses, tuning for a standard APE unit should move backwards in time to the start of the sequence the same way it classically does in the scalar RPE model. Instead, we would expect this heterogeneity to reflect feature-specific computation within each APE circuit (**Fig. 8f**).

To assess whether a feature-specific APE model could account for the results in Jin and Costa^5^, we simulated a simplified version of their task (**Fig. 8e**) by augmenting the choice task from **Fig. 8a** with a requirement of 3 presses of the lever to receive the reward (see Methods). To create predictions for DA responses on this task, we used both the standard APE model and a feature-specific APE model (**Fig. 8f**) that functions like the feature-specific PE model but uses the state features to produce a set of prediction errors for (e.g.) left presses as outcomes instead of rewards (see Methods). For simplicity, we only analyze the left-selective circuits within each of these, but we note for completeness that they both include parallel left- and right-selective circuits.

For both models, the state representation was a one-hot encoding of the press count (**Extended Data Fig. 7b**). In the standard APE model, this predictably resulted in a response that peaked at the first press for all neurons and did not produce specific press selectivity (**Fig. 8g, right**). In contrast, the feature-specific APE model showed uniform selectivity for press counts across all DA neurons (**Fig. 8g, left**), with each DA neuron reflecting the tuning of its upstream state neuron. Thus, the feature-specific APE model recapitulates the experimentally reported press-selectivity across the population^5^. Furthermore, each feature-specific DA neuron exhibited a transient decrease in activity after its preferred press due to the feature-difference term in Equation 1, which is a property that has been observed in SNc DA neurons^5,21,67–69^.

One prediction of our model is that DA heterogeneity should be directly tied to the upstream cortical/striatal state features. For example, Jin and Costa^5^ observed that from day 1 to day 12 of training, the proportion of first-press and last-press neurons increased in tandem for both SNc DA neurons and striatal MSNs, with first-press increasing the most. To the extent the population of DA responses is indeed coupled to the upstream code, our model predicts that these changes imply a remodeling of the upstream population (simulated in **Fig. 8h** by changing the state representation in our model of the task so that the first and last press are overrepresented; as expected, this induces the corresponding representational shift at the DA level). Indeed, this is consistent with evidence (including from Jin and Costa^5^ and also from other groups and tasks) that overtraining of this sort is also accompanied by a “chunking” of corticostriatal representation of the action sequence to “chunks” so that the overall sequence becomes identified with its starting and concluding actions^70,71^.

## Discussion

Here we propose a new theory which helps to reconcile recent empirical reports of DA response heterogeneity to the classic idea of DA neurons as encoding RPEs. Most previous work has considered “outcome-specific” PEs that describe heterogeneity between parallel circuits for predicting different outcomes. We introduce a complementary “feature-specific” model, where heterogeneity arises from the high-dimensional state input to each such circuit. We show how this model, but not alternative models based on outcome-specific PEs, produces heterogeneous responses to task variables, but relatively uniform responses to reward, recapitulating recent empirical work^4,20^. We also test the model’s prediction that heterogeneous DA responses are not simply responses to sensory and behavioral features of the task, but instead reflect components of the RPE with respect to a subset of the features. Finally, we show how the principles of outcome- and feature-specific PEs combine to explain different aspects of movement-related heterogeneity in SNc DA neurons^5^.

### Feature-specific vs outcome-specific PEs: Physiology

Our central observation is that TD models imply two distinct candidate sources of heterogeneity across DA units: that arising from the target outcome (e.g., reward) and that arising from the state input (e.g., cues and other task features). Most previous theories have explained DAergic heterogeneity by positing multiple distinct error signals, most often each specialized for learning to predict a different outcome^31–33,35–39,58,72–76^. For instance, a family of error signals could be used to learn to predict different outcomes such as rewards versus punishments^31^, to learn the rewards associated with different actions or effectors^40–42,58^, to predict rewards at different temporal scales^26,38^, or to track different goals and subgoals in a hierarchical task^74,75,77^. In contrast, the basic insight of our new model is that within any such outcome-predicting circuit, nonuniform input projections from an arbitrary population code for state can give rise to diverse patterns of simultaneous, multiplexed responses to different nonreward task events. Collectively, these responses constitute a population code over feature-specific PEs in an otherwise standard TD learning setting aiming (in each circuit) to predict each single, scalar outcome like reward.

Importantly, both feature-specific and outcome-specific models are compatible and likely coexist, capturing different aspects of DA heterogeneity. We can differentiate their respective contributions to observed heterogeneity in a number of ways. The first is that they imply different relationships between heterogeneous responses at outcome (to rewards, omissions, or punishments^15,17,21,24,30,33,78–83^) versus heterogeneous responses to other task events, including stimuli and movements^4,5,7,13,20,21,23,27,67,82–85^. Outcome-specific PEs imply that variation between neurons in their responses to stimuli ultimately arises from those neurons specializing in different outcomes. For this reason, these models fail to capture the coexistence in VTA DA between heterogeneous responses to task variables alongside a more uniform main effect of reward^4^. This is instead a hallmark of a feature-specific RPE, in which different neurons represent different stimulus-specific components of a PE for a common reward. Our simulations show that different outcome-specific models can either produce heterogenous coding of cues and task variables by predicting many outcomes, but then fail to respond to reward (**Fig. 6**, successor representation) or instead maintain the overall reward effect by focusing on predicting a narrower range of reward-related outcomes, but with limited variability in responses to cues and other task variables (**Fig. 7**, distributional RL).

A related prediction of the feature-specific RPE is that DA encoding of task features, in general, actually reflects components of RPE with respect to each feature, rather than strictly the main effect of each feature itself. For instance, in our data and in the feature-specific RPE model, different DA neurons are tuned for different cues (left vs. right), but each population further distinguishes confirmatory from disconfirmatory cues (**Fig. 4b,c**), which differ in their reward consequences. In contrast, different SR neurons respond to the cues themselves, not their reward associations (**Fig. 6d,e**). An important aspect of the data is that the responses are tied to individual features – consistent with the feature-specific model’s unique claim that different neurons represent RPE components for left cues and right cues separately. This stands in contrast to a distributional RL model (**Fig. 7a**) and also to classic scalar TD (**Fig. 1a**), both of which also predict that cue responses should be modulated by their reward associations (e.g., confirmatory vs. disconfirmatory **Fig. 7e**), but wrongly predict that all neurons should respond similarly to left vs. right cues, which do not have distinct reward associations (**Fig. 7d**).

Importantly, because both frameworks are so general – different PEs can be defined for any vector of outcomes, and also for any vector of features – we have here emphasized qualitative predictions that aim to capture what is common to each entire family^86^. We have also, for concreteness, used a deep network model to learn a detailed and reasonably realistic high-dimensional feature set with which to simulate the model and expose its key behaviors. However, this model involves a number of simplifications (for instance, it does not vary running speed or other kinematics, and does not carry any information from trial to trial) which mean that it cannot in principle capture some types of responses reported in the neural data^4^, and is also likely to quantitatively differ with respect to others (e.g. head direction, which is more constrained in simulation). One aspect of the response that is notably different in simulation is position-related ramps. While these do arise in simulation – and demonstrate how feature-specific PEs can, for appropriate features, capture even temporally extended responses – they tend in the model to occur only over shorter distances than in DA recordings, possibly reflecting how the model features were built (unrealistically) via backpropagation through time. In any case, the phenomenon of positional ramping in DA is under active investigation, and a full explanation is likely to involve a number of additional considerations (e.g. uncertainty) not included in the current model^87–89^.

### Feature-specific vs outcome-specific PEs: Anatomy

An important dimension predicted to distinguish feature-from outcome-specific heterogeneity is anatomical. DA neurons reporting different outcome-specific PEs should be linked to their targets in a closed loop, meaning this form of heterogeneity should be found across projection-defined DA populations. In contrast, heterogeneity arising from the feature-specific PE model is nested within each outcome-specific circuit, meaning it should exist even within a projection-defined population of DA neurons. In fact, feature-specific decompositions of a PE will best retain the strengths of the scalar model when units representing different aspects of PE have overlapping targets, because this would allow PEs for different aspects of state to mix into a common update.

Together, these considerations lead to the overall prediction that the characteristics of outcome-vs feature-specific PE variation should predominate, respectively, between vs within different projection-defined DA populations. Formally investigating this prediction (e.g., by systematically comparing cue and outcome responding across different projection-defined DA populations) is the key open empirical test of our framework. However, existing data are generally consistent with the hypothesis that the sorts of heterogeneity associated with outcome-specific differences tend to arise between distinct DA nuclei or target regions^7,90,91^, in contrast to the hallmarks of feature-specific PEs we study here, which occur within VTA^4,20^ and also within SNc^5^. Thus, in SNc and DMS, compared to VTA and NAc, movement responses tend to emerge while reward responses decline (a combination consistent with outcome-rather than feature-specific variation; **Fig. 8**)^5,7,21,23,85^; DA neurons responding to threat rather than reward have been reported in projections from lateral SNc to the tail of the striatum^30,92^; and recent data suggest that the differential responsivity to RPE magnitudes that is characteristic of distributional RL also emerges across DA projections to distinct striatal subregions^90,91^. A related point is that, since the feature-specific PE model predicts that averaging over features will recover the original scalar PE, the types of variation it envisions will tend to be washed out by methods like photometry that involve averaging over many neurons.

While more anatomical data is needed to fully clarify the scope of the feature-specific and outcome-specific models, existing data regarding inputs to DA neurons, as well as DA innervation to striatum, already provides important information. Consistent with our feature-specific model, inputs to DA neurons (especially from striatum), which in our model represent the state representation, are topographic, and plausibly more precisely so than the ascending DA projection^44,45,48,49^. These afferent neurons are known to reflect the same sorts of heterogeneity with respect to task features as we consider at the DA level^5,61–66^, consistent with our framework in which the set of DA units that collectively target some striatal domain are assumed to vary in their feature inputs (e.g. **Fig. 8h**). Regarding DA projections, individual DA neurons branch extensively to innervate large areas (∼1 mm^3^) of striatum^11^, and volumetric propagation of released DA to extrasynaptic DA receptors may further blur DA’s effect^9,93–95^. This suggests a limited number of sufficiently non-overlapping outcome-specific PE circuits are possible, although how many remains unknown. Recent work partly moderates these concerns by emphasizing that DA signaling may be somewhat more precise than the initial observations suggest, in that only a minority of DA varicosities have active release sites, and DA reuptake can limit diffusion^96–98^.

### Action prediction error

In extending our feature-specific PE model to SNc/DMS DA movement signals (Figure 8), we made use of the action prediction error (APE) model^40,41^, which is the hypothesis that dopamine neurons can represent a PE computed for prediction of future actions. We showed how fusing this idea with our feature-specific RPE model to create a feature-specific APE model can, in a unified framework, explain features of movement signals in DA neurons, such as the pattern and heterogeneity of action responses, and the close relationship between striatal action representations and SNc DA representations^5^ (Figure 8).

However, we stress that evidentiary support for the existence of a DA APE in the brain is not yet definitive. Though recent work has shown that DA in the tail of the striatum elicited by cue-evoked action execution decreases over time ^40^, supporting the idea that action responses are modulated by predictability rather than simple motor correlates, other experiments must still be conducted in order to establish that DA neurons encode an APE fully analogous to the TD RPE. Notably, a TD prediction error transfers with learning from the predicted outcome to cues predicting it (at least depending on the time discount parameter, which might also vary between circuits^26,99,100^). By analogy, a TD APE predicts that cues that elicit actions should produce DA signals that increase with training, and that “omission” of an action predicted by a cue (say, by introduction of a no-go signal) should yield a reduction in DA. Another primary, testable prediction of the APE includes that action expectation should be graded and dynamic – that is, tasks in which cues predict actions only probabilistically should induce dopaminergic responses to both cues and actions whose magnitudes smoothly vary with the response probability, and track it dynamically during learning. To our knowledge, these predictions have not yet been tested and reported in the literature.

Furthermore, while the APE idea helps to reconcile DA action responses with the RPE framework, it is not the only example of an outcome-specific PE for which evidence is emerging. For instance, other recent work on DA neurons projecting to the tail of the striatum identifies responses to threat or novelty (again modeled with an outcome-specific PE for these variables) that decline with learning and are associated with outcome-specific unconditioned and learned responses such as avoidance^30,34^. More broadly, the overall question of how large-scale DA circuits are functionally and anatomically organized remains a major open question in the field; we believe that introducing a distinction between outcome-based heterogeneity and feature-based heterogeneity, nested at different spatial scales, may present a starting point for organizing that discourse.

### Credit assignment and outcome- vs feature-specific PEs

Previous outcome-specific PE models have typically been motivated as enabling enhanced computational function: additional PEs compute additional quantities^31–39,58^. In contrast, the motivation for our model is a pair of more practical issues with this picture. The first is the need for distinct closed-loop circuits for each PE, which likely constrains the scale of heterogeneity that can play this role. The second is the requirement that information about many widely dispersed state features be brought together as inputs for value prediction and RPE. We suggest that these two issues are related: while it is likely inevitable that inhomogeneous state feature input drives response variation within a projection-defined DA population, the same overlapping ascending projections that constrain the outcome-specific PEs also help to promote convergence across these feature-specific PEs.

Such convergence, in turn, addresses a central computational function: credit assignment over multidimensional feature spaces. Normatively, the scalar RPE arises algebraically from gradient descent and also from statistical inference, when a prediction is made from a weighted sum of features (i.e., the delta rule is a special case of both backpropagation and also Bayes’ rule^101–103^). Functionally, the effect of such a common RPE is to enable stimuli to compete with one another to explain observed rewards, each stimulus learning uniquely to explain what cannot be explained by the others. In Pavlovian conditioning, this insight goes back to the classic model of Rescorla and Wagner, who pointed out that such competitive credit assignment via a shared RPE is observable behaviorally in the blocking effect: an animal will not learn that some stimulus predicts reward, if the reward is already predicted on the basis of another stimulus^104,105^.

These considerations provide a normative, computational rationale why – to whatever extent RPE heterogeneity arises from feature inputs rather than outcome targets – it is advantageous that they mix (as in **Fig. 1c**) rather than remain separate (as in **Fig. 1b**), and thereby reveal an unrecognized virtue of the diffuse ascending anatomy. They also offer, via the blocking effect, a behavioral test of this prediction. If stimuli instead have separate errors, PE-driven learning still works for each stimulus individually (this was the basis of conditioning models predominant prior to Rescorla-Wagner^106,107^), but they will not engage in competitive credit assignment. This means our model degrades gracefully if, in practice, mixing over the feature-specific RPEs is imperfect (e.g., if ascending projections do not perfectly overlap). To whatever extent different DA subpopulations represent RPEs specific to different features, but without anatomically overlapping projections, their respective cues will fail to block one another behaviorally. For instance, if VTA DA responses to towers are largely lateralized in opposite hemispheres in Pavlovian conditioning, as they are in the Engelhard et al^4^ task, then (although the data lack projection tracing) their targets are unlikely to overlap, and such a failure of blocking could be expected between left and right towers.

Although cue competition via a common error signal offers a well-justified and simple approach to credit assignment, if cues are instead separated in a parallel-circuit architecture like the outcome-specific PE models, then credit assignment requires an additional computational approach. Such models, collectively known as “mixtures of experts,” use separate prediction modules associated with different subsets of stimulus features and represented in different closed-loop channels^25,108,109^. Mixture of expert schemes offer more individualized control, to “divide and conquer” and focus learning on context-relevant dimensions. However, unlike cue competition through a scalar error signal, in this case credit assignment requires an external arbitration mechanism to determine which modules to rely on in any particular context^25,108,109^. One recent model^25^ uses this type of architecture to suggest a different set of mechanisms addressing some aspects of the problems we discuss here. In particular, this work suggests that even within a broadcast error signal, the timing of a scalar DA response (arriving earlier or later due to wave-like response propagation) can target the signal to different modules. The study also suggests arbitration principles for externally assigning credit to different modules by routing waves depending on circumstances, though the detailed neural basis of such arbitration remains to be explored. Overall, to what extent the brain relies on either credit assignment mechanism remains a question, but the present framework clarifies how these issues could be examined via behavioral (e.g. blocking), projection-tracing and neurophysiological studies.

### Population codes for state

Although much RL modeling in neuroscience assumes a simple state representation that is hand-constructed by the modeler^2,3^, in general, an effective state representation depends on the task, and the brain must learn or construct it autonomously as part of solving the full RL problem. How it does this is arguably the major open question in these models. Indeed, in AI, recent progress on this problem (notably using deep neural networks) has been the main innovation fueling impressive advances scaling up otherwise standard RL algorithms to solve realistic, high-dimensional tasks like video games^55,110^. In psychology and neuroscience also, there have been a number of recent theoretical hypotheses addressing how the brain might build states, such as the successor representation and latent-state inference models^51,111–115^. But there exist relatively few experimental results to assess or constrain these ideas, for instance because learning behavior alone is relatively uninformative about state, whereas in the brain it is unclear which neural representations directly play this role for RL (e.g., grid cells vs. place cells for spatial tasks^114,116^).

A main consequence of the feature-specific PE model is that, if it is correct, then the heterogeneous DAergic population itself gives a new experimental window, from the RL system’s perspective, into the brain’s population code over state features. While the model itself is agnostic to the feature set used, the various DA responses should in any case reflect it. This builds on previous work showing that even a scalar DAergic TD error signal can be revealing about the upstream state that drives it^50,52,53^, but on the new theory, the feature-specific vector DA code much more directly reflects the upstream distributed code for state. Thus, the theory offers a general framework to reason quantitatively about population codes for state. This should enable new experiments and data analyses to infer the brain’s specific state representation from neural recordings and in particular to test ideas about how it is built: how it changes across different tasks and as tasks are acquired.

## Acknowledgements

We thank Alvaro Luna and Juan Lopez for their help with the VR software system, Will Dabney for discussion on the distributional RL model and providing the imputation function for the distributional RL model, Michelle Lee and Erin Grant for help with training the deep RL network, Peter Dayan, Ari Kahn, and Lindsey Brown for comments on this work, and additionally the BrainCOGs team and the entire Daw and Witten labs for their help. This work was supported by an NSF GRFP (RSL), 1K99MH122657 (BE), NIH R01 DA047869 (IBW), U19 NS104648-01 (IBW), ARO W911NF-16-1-0474 (NDD), ARO W911NF1710554 (IBW), Brain Research Foundation (IBW), Simons Collaboration on the Global Brain (IBW), and the New York Stem Cell Foundation (IBW).

## Figures

**Extended Data Figure 1:**
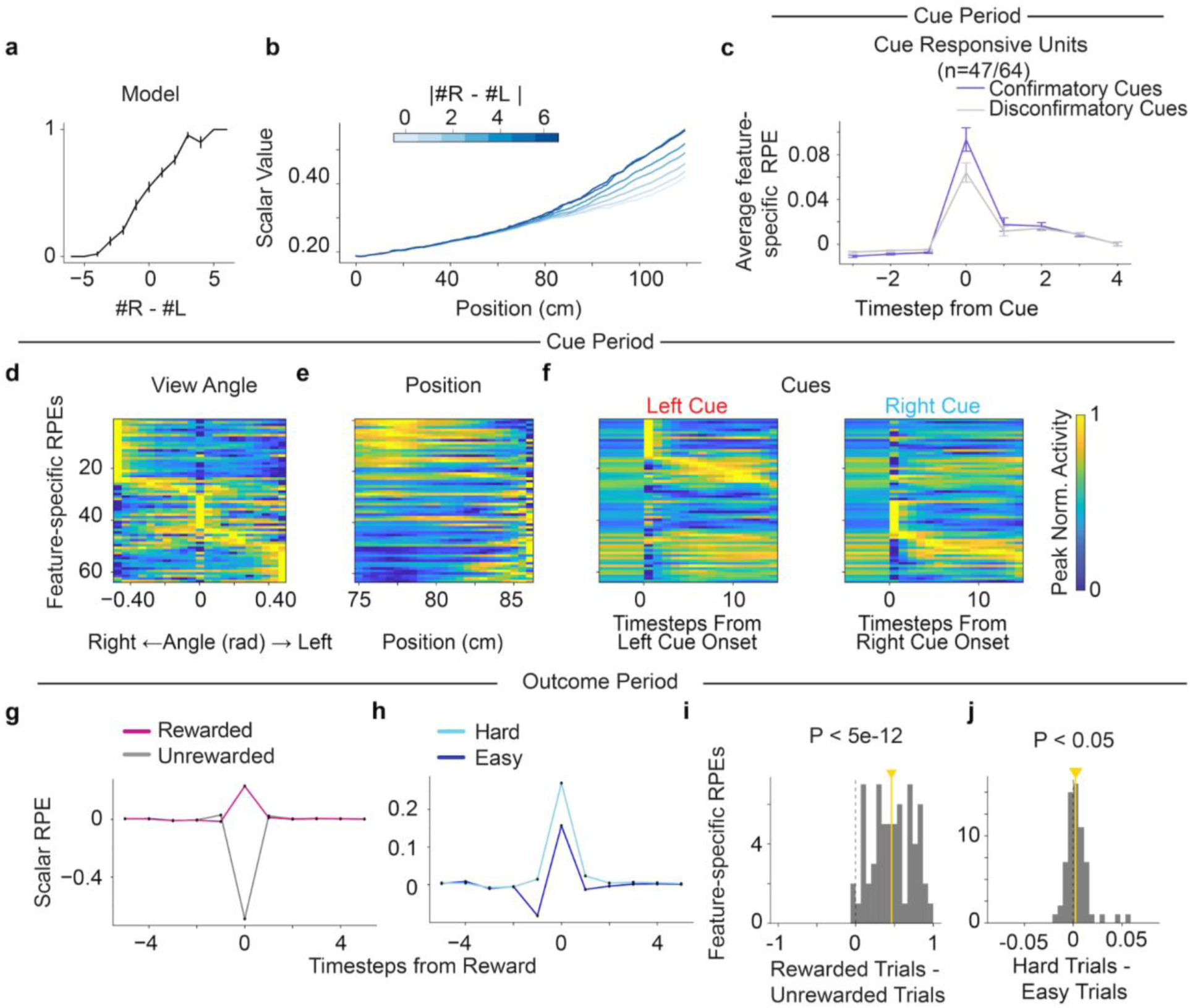
Deep reinforcement learning network retrained with the same task and a different seed. **(a)** Psychometric curve showing the retrained agent’s performance. **(b)** Retrained agent’s scalar value during the cue period decreased as a function of trial difficulty (defined as the absolute tower difference, blue gradient). **(c)** Retrained agent’s feature-specific RPE units’ response to confirmatory (purple) and disconfirmatory (gray) cues. Response is averaged across cue-sensitive units only (N = 47/64). **(d)** Retrained agent’s feature-specific RPE units averaged across trials plotted with respect to view angle. Each row represents one unit’s peak normalized response to the view angle. **(e-f)** Same as **(d)** but for the agent’s **(e)** position and **(f)** left (red) and right (blue) cues. **(g)** Retrained agent’s scalar RPE time-locked at reward time for rewarded (magenta) and unrewarded trials (gray). **(h)** Same as **(g)** but for rewarded trials with different trial difficulties, with hard trials (light blue) and easy trials (dark blue) defined like in Fig. 5d. **(i)** Histogram of retrained agent’s feature-specific RPE units’ response to reward minus omission at reward time (P < 5e-12 for two sided Wilcoxon signed rank test, N = 64). Yellow line indicates median. **(j)** Same as **(i)** but for rewarded trials plotted against trial difficulty (P < 0.05 for two sided Wilcoxon signed rank test, N = 64). In **(j),** there is an outlier data point at 0.16 for a feature-specific RPE unit showing strong reward expectation modulation.

**Extended Data Figure 2:**
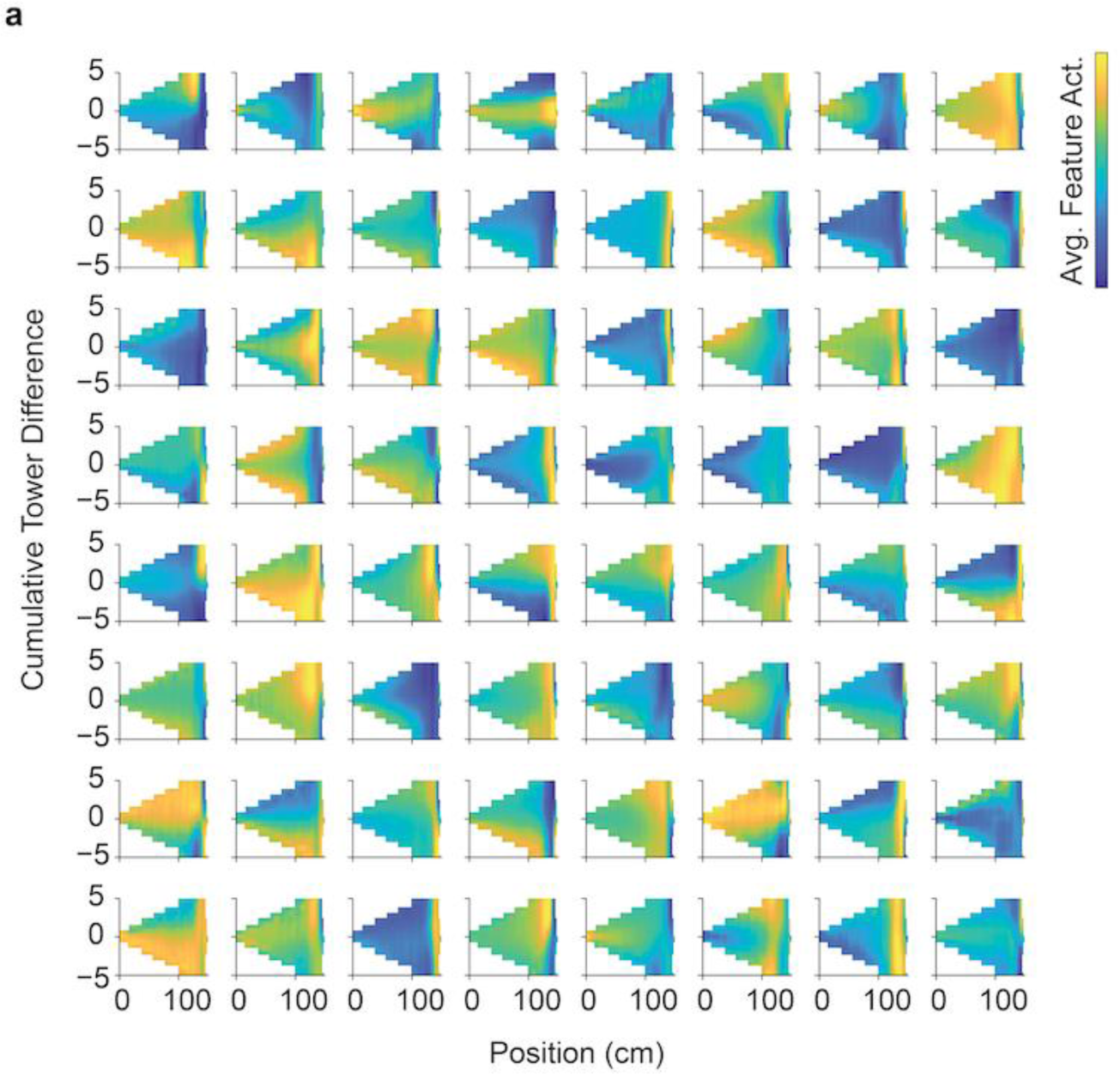
Tuning of 64 LSTM feature units to position and evidence. **(a)** Each panel shows an individual feature unit and how it is tuned to the agent’s position in the maze and the cumulative tower difference at that position.

**Extended Data Figure 3:**
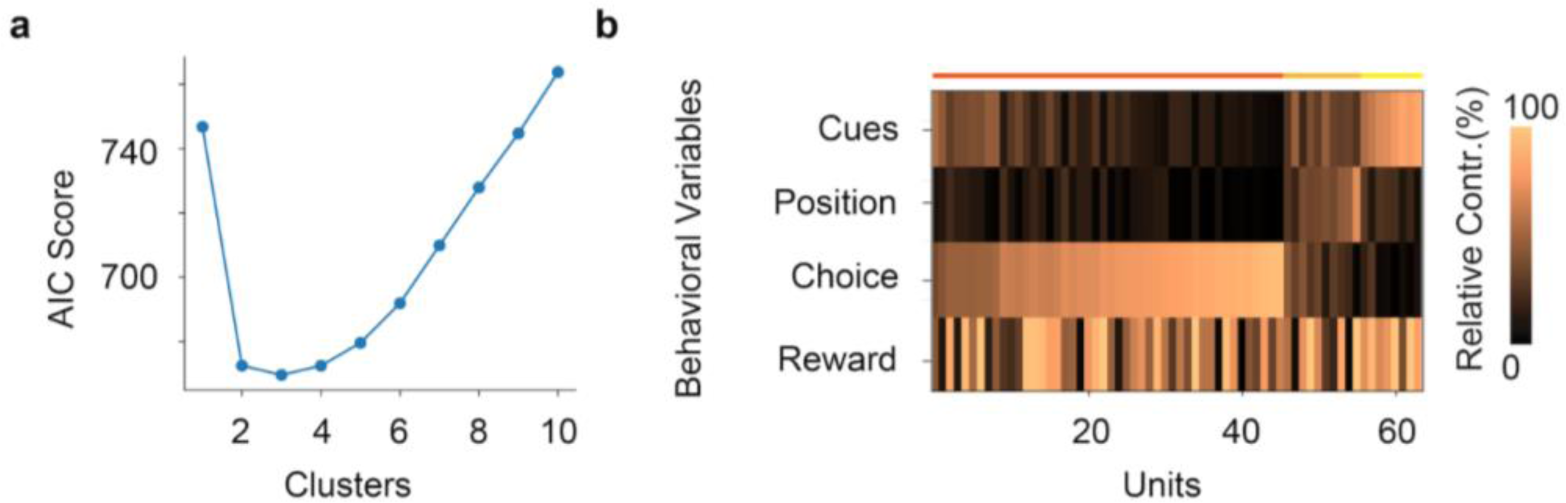
Feature-specific RPEs clustered to behavioral features of the task to match Engelhard et al^4^ clustering analysis. **(a)** Optimal number of clusters was 3, selected by minimizing the AIC scores for models with different numbers of clusters. **(b)** Relative contribution of the behavioral features including cues, position, choice and reward response for the 64 units sorted based on the highest probability belonging to the cluster, with colored lines on top indicating the cluster identity. Relative contribution is defined as the percentage of the explained variance for the partial model not including the variable versus the full model.

**Extended Data Figure 4:**
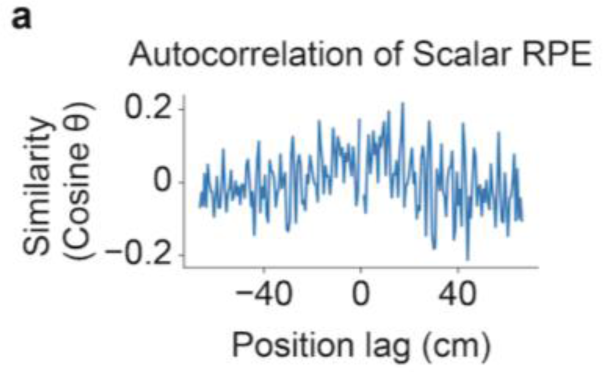
Scalar RPE signal does not reflect the incidental high-dimensional features. **(a)** Scalar RPE signal autocorrelated across time (similarity defined as the cosine of position-lagged scalar RPE responses) does not show peaks at position lag 43 cm for the wall-pattern repetition location.

**Extended Data Figure 5:**
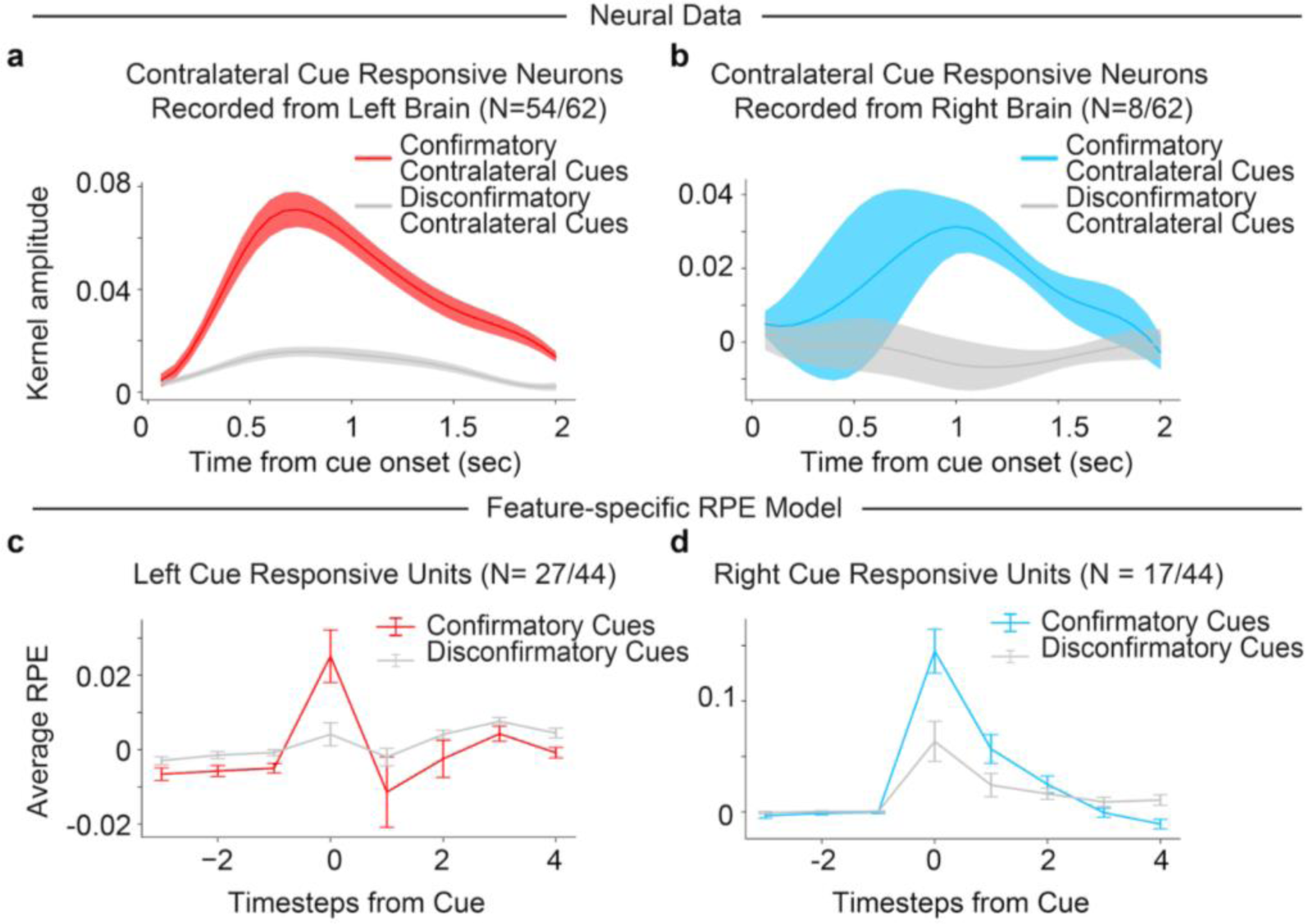
Cue responses in model and DAergic neurons represent a vector code of lateralized response and RPE. **(a)** Average responses of the contralateral cue responsive DA neurons^4^ only recorded from the left hemisphere (N= 54/62 neurons, subset of contralateral cue responsive neurons from Fig. 4c) for confirmatory (red) and disconfirmatory (gray) contralateral cues. Colored fringes represent ±1 s.e.m. for kernel amplitudes. **(b)** Same as **(a)** but for DA neurons recorded on the right hemisphere (N = 8/62) responding to confirmatory (blue) and disconfirmatory (gray) confirmatory cues. **(c-d)** Same as **(a-b)**, but for feature-specific RPE model units responding to **(c)** left cues specifically (N = 27/44, subset of the cue responsive neurons from Fig. 4b that were modulated by left cues only) and **(d)** right cues specifically (N = 17/44). Error bars indicate ±1 s.e.m.

**Extended Data Figure 6:**
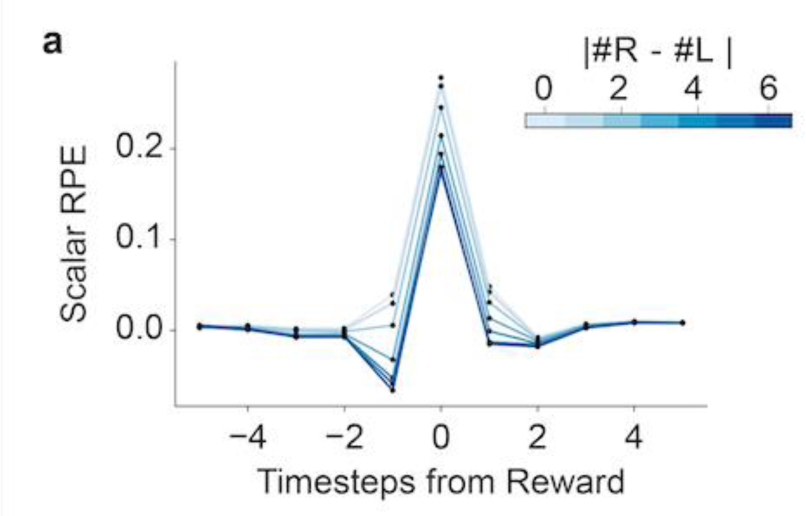
Scalar RPE shows fine-grained reward expectation modulation. **(a)** Scalar RPE’s response modulated by reward expectation given by the difficulty of the task, defined as the absolute value of the final tower difference (blue gradient) of the trial.

**Extended Data Figure 7:**
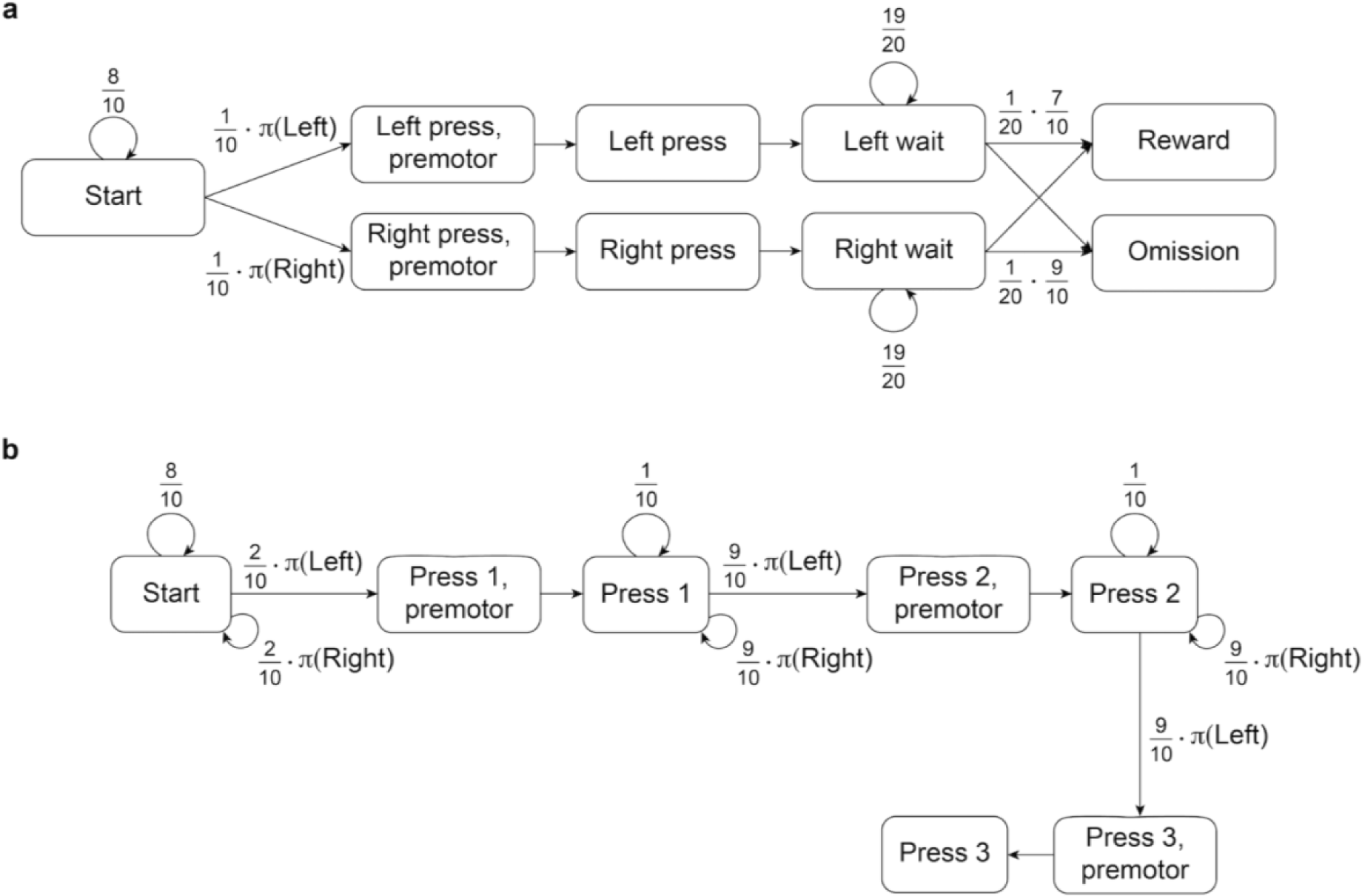
State diagrams for simulations of the Parker et al^7^ and Jin and Costa^5^ tasks in Fig. 8. **(a)** State diagram for the Parker et al^7^ task simulation. **(b)** State diagram for the Jin and Costa^5^ task simulation. In both panels, arrows indicate probabilistic transitions between states, with probabilities described by the arrow labels. Unlabeled arrows denote transitions with the remaining probability to make the total sum to 1. *π*(*x*) refers to the probability of executing action *x* under the agent’s behavioral policy *π*.

## Methods

### Behavioral Task

#### Simulations

At every trial, the agent was placed at the start of a virtual T-maze, with cues randomly appearing on either side of the stem of the T-maze as the agent moved down the maze. On each trial, one side was randomly determined to be correct, and the number of cues on each side was then sampled from a truncated Poisson distribution, with a mean of 2.29 cues on the correct side and 0.69 cues on the incorrect side. In order to prevent the agent from forming a side bias, we used a debiasing algorithm to ensure that the identity of the high probability side changed if the agent kept choosing one side^117^. To match the procedure from the mouse experiment, we also oversampled easy trials (trials in which only one side had 6 total cues, and the other had no cues) by ensuring that they were 5% of the trials. The agent moved down the maze at a constant speed of 0.638 cm per timestep, and could also modulate its view angle with two discrete actions corresponding to left and right rotation. (A third discrete action moved forward without changing the view angle.) The cue region was 85 cm, and the cues were placed randomly along the cue region under a uniform distribution, but with the restriction that cues on either side had a minimal spatial distance of 14 cm between them. Each cue first appeared when the agent was 10 cm from the cue location and disappeared once the agent passed the cue by 4cm. After the cue region, there was a short, 5 cm delay region before the agent’s final left or right action determined their choice of entering either arm in the T-maze. If the agent turned to the arm on the side where more cues appeared, they received a reward. The agent was also given a sensory input in the model indicating whether it made the correct or wrong turn.

#### Neural data

The task simulated above was a streamlined version of that used in the mouse recordings in Engelhard et al.^4^ In particular, the rules for spacing and visual appearance of cues were the same, but the simulated controls were simplified (to discrete actions) and the maze was shorter (to facilitate neural network training). Thus the mice could control their speed and direction of movement more continuously by running on a trackball, and in this way traversed a maze that had a 30 cm start region (with no cues), a 220 cm cue region, and an 80cm delay region before the T maze arms. The mean numbers of cues were correspondingly larger: 6.4 on the correct side and 1.3 on the incorrect side. At reward time, the mice received a water reward if they made the correct choice; if a mouse made an incorrect choice, it was given a pulsing 6-12 kHz tone for 1 second. Before the next trial, the virtual reality screen froze for 1 second during reward delivery, and blacked out for 2 seconds if the mouse was rewarded or 5 seconds if the mouse failed.

### Virtual Reality System and Deep Reinforcement Learning Model

A deep reinforcement learning network was trained on the evidence accumulation task. As input, the network took in 68 by 120 pixel video frames in grayscale. The model had 3 convolution layers to analyze the visual input, an LSTM layer to allow for memory, and output to 3 action units and 1 value unit. The first convolutional layer had 64 filters, a filter size of 8 pixels, and a stride of 2 pixels; the second convolutional layer had 32 filters, a filter size of 2 pixels, and a stride of 1 pixel; the third convolutional layer had 64 filters, a filter size of 3 pixels, and a stride of 2 pixels. The convolution layers fed into a fully connected layer, which fed into the LSTM layer with 64 units along with a second input, a one-hot vector of length 2 which flagged whether or not the agent was rewarded the end of the trial. The reward input into the LSTM was meant to replicate the sensory input that the mouse experienced when it was rewarded with water or received a tone for failing the trial. The hyperparameters for the convolutional layers were optimized with a grid search of various filter numbers and sizes trained on supervised learning for recognizing towers. The convolution layer feeds into a fully connected feature layer of 128 units, then to 64 LSTM feature units, 64 value units, and 64 feature-specific RPE units.

We use the same MATLAB virtual reality program (ViRMEn software engine^118^) from the original neural recordings^4^, which we altered to accommodate the agent’s movement choices of forward, left, and right. While in the stem of the T-maze, the agent always moved forward at a constant rate per timestep. The constant speed ensured that at every trial, the agent always took the same number of timesteps to traverse the stem of the T-maze. The agent could choose to rotate left or right, which would alter the view angle 0.05 rads up to the limit of -π/6 and π/6 rads. The agent could also choose to move forward without changing their view angle. After the delay region in the T-maze, the agent’s left and right movement would no longer alter its view angle, but instead determined which arm the agent chose.

In order for the deep RL agent to interact with the ViRMEn software, we created a custom gym environment using OpenAIGym’s *gym* interface^119^. Our custom virtual reality gym environment defined the forward, left, and right movements the agent could make, and sent in the movement choices to the ViRMEn software which in turn returned updated video frames.

We trained the network to maximize obtained reward using the Stable Baselines^120^ (version 2.10.1) implementation of the Advantage Actor Critic (A2C) algorithm^56^. The loss function for training using the A2C algorithm with weight parameters *θ*, action at time t *a*_*t*_, state *s*_*t*_ and reward *r*_*t*_ is given as:

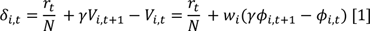

The first term is the actor loss, given as the log of the policy *π* multiplied by a sample of the 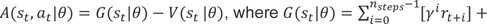 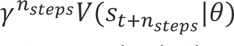 is an n - (defined as *n*_*steps*_ in the model) step Bellman estimator for the return and *V* is the value (critic) function. The second term is the critic loss, or the squared error of the value function estimate with respect to the return *G*. The final term is an entropy term which regularizes the policy to increase exploration. *β* and *η* are hyperparameters (listed below) to trade-off the effects of the regularization and the losses.

All hyperparameters used for the deep RL agent can be found in Table 1. We trained the model until it reached a performance of 80% or higher correct choices, which took 140.8 million timesteps or approximately 900,000 trials. To show that the results were consistent across multiple starting weights, we retrained the agent with fresh random weights and found that the new features replicated the main results of the paper (**Extended Data Fig. 1**).

**Table 1:** Hyperparameters for A2C Algorithm.

| Parameter (Variable Name in Stable Baselines) |  |
| --- | --- |
| Number of Parallel Environments ($n_{env}$) | 8 |
| Number of steps until each environment updates ($n_{steps}$) | 140 |
| Discount Factor (gamma $\gamma$) | 0.99 |
| Learning Rate | 0.0025 |
| Value function coefficient for loss calculation (vf_coef $\beta$) | 0.25 |
| Entropy coefficient for loss calculation (ent_coef $\eta$) | 0.01 |
| Maximum value for gradient clipping (max_grad_norm) | 0.5 |
| RMSProp decay parameter (alpha) | 0.99 |
| RMSProp momentum parameter (momentum) | 0.0 |
| RMSProp epsilon (epsilon) | 1e-5 |

We also ran a few additional simulations to learn more about the training process. First, pretraining the convolutional layers of the network to recognize towers significantly reduced training time, as well as shortening the delay region from the final tower appearance to the final tower location. The agent was also sensitive to the *n*_*steps*_ hyperparameter, which determined the batch gradient size; specifically, a shorter *n*_*steps*_ early in training helped with the agent learning the task above chance, but longer *n*_*steps*_ was needed to master the task to reach the performance we see in the paper. We also trained the network with only 8 LSTM units, but the agent could not learn the task even after 200 million timesteps. A larger and overparameterized network was evidently needed for the agent to understand the task and keep track of the towers seen, even though the theoretical state space sufficient for the task is three dimensional (left tower count, right tower count, and position of the agent).

After training, we froze the weights and took the final output layer of the network before the action and value units (i.e., the LSTM output) as the features for vector value and feature-specific RPE (**Equation 1**, **Fig. 2b**). (Note that this just corresponds to decomposing the scalar value and RPE units in the original A2C network into vectors, with one value component for every LSTM-to-value weight, and a corresponding RPE component: i.e. the feature-specific RPE model, being algebraically equivalent to TD, is just a more detailed view of the A2C critic). In this way, we calculated the feature-specific RPE at every point in the trial, for a total of 5000 simulated trials. We defined the outcome period to be the 5 timesteps before and after the reward. We defined the cue period as the first 140 timesteps of the maze, which occurred at the same positions on every trial since the agent always moved forward the same amount at each timestep.

### Neural Data

This article analyzes data originally reported in Engelhard et al^4^, the methods of which we briefly summarize here and below^4^. We primarily re-analyzed the neural recordings during the virtual-reality experiments, in which we used male DAT::cre mice (n=14, Jackson Laboratory strain 006660) and male mice that are the cross of DAT::cre mice and GCaMP6f reporter line Ai148 (n=6 Ai148xDAT::cre Jackson Laboratory strain 030328).

VTA DA neurons were imaged at 30 Hz using a custom built, virtual-reality compatible two photon microscope equipped with pulsed Ti:sapphire laser (Chameleon Vision, Coherent) that was tuned to 920 nm. After imaging, we removed trials in which mice were not engaged in the task, primarily those found close to the end of the session when animal performance typically decreased. Average performance across sessions on all trials was 77.6 +/- 0.9% after removing trials (compared to 73.3 +/- 1.1% including all trials). Ultimately, we used 23 sessions from 20 mice (one session per imaging field, each session with at least 100 trials and minimal performance of 65%).

After preprocessing the imaging data, we performed motion correction procedures to eliminate spatially uniform motion and spatially non-uniform, slow drifts. The dF/F was derived by subtracting the scaled version of the annulus fluorescence from the raw trace (correction factor of 0.58) and smoothed using a zero-phased filter with 25 point center Gaussian with 1.5 sample points standard deviation. We then divided dF/F by the eighth percentile of the smoothed and neuropil-corrected trace based on the preceding 60s of recording. After examining the dF/F, we only included neurons that were stable for at least 50 trials. The full dataset we used for renalaysis has 303 neurons spread across 23 sessions from 20 mice.

### Cue Period Responses

For the heatmaps in **Fig. 2e-j**, each row represents a feature-specific RPE unit or neuron’s response to the behavioral variable rescaled so that their maximum value is at 1 and minimum value is at 0. The example plots above show the unnormalized response. For the neural data, we used the same encoding model from our previous work^4^ to predict neural activity with a linear regression based on predictors including cues, accuracy, previous reward, position, and kinematics, to isolate temporal kernels that reflect the response to each cue. The code for this encoding model can be found at https://github.com/benengx/encodingmodel. To determine which neurons are displayed for the heatmaps in **Fig. 2h-j**, we used the same criterion as in Engelhard et al^4^, which was to include neurons with a statistically significant contribution of that behavioral variable in the full encoding model relative to a reduced model, based on an F-test (P = 0.01), with comparison to null distributions produced by randomly shifted data to account for slow drift in the data.

### Encoding Model and Clustering Analysis

For **Extended Data Fig. 3**, we adapted the encoding and clustering models from Engelhard et al^4^ to estimate the relative contributions of various behavioral cues and variables to the model’s simulated DA responses, and then cluster them using a Gaussian mixture model. First, we used multiple linear regression to model cue period activity, per simulated neuron, as a function of three behavioral features: cues, position, and the agent’s choice. Each feature was coded with a group of regressors; we used the regression to measure the relative contribution (defined in terms of variance explained) of each group to each neuron’s responses, for clustering (below). Coding of the variables followed that used for dopamine neurons in Engelhard et al^4^: specifically, the cue variables were coded as event-locked timeseries, with the appearance of left and right cues convolved with an 8 degrees of freedom regression spline basis across 10 timesteps of duration. Position was entered as a degree-3 polynomial series on the continuous variable, (i.e. we included the first, second, and third powers of position). We then chose the optimal number of predictors using a fivefold cross-validation over trials. The agent’s final choice was coded with three variables as one-hot vectors, indicating the decision time for a left choice, and one and two steps before that. (There were no separate variables for choosing the right arm since we also included a bias variable in the regression.)

For the outcome period, we modeled timesteps following reward using two sets of regressors, representing (by one-hot event-triggered lagged indicators) the time of reward (on rewarded trials only) and time of outcome (comprising the time of either nonreward or reward). Each comprised 7 one-hot vectors indicating the event time (reward and/or nonreward) time and the 6 timesteps following. No bias variable was included.

After regressing cue and outcome period activity, the relative contribution and GMM clustering procedures were performed as in Engelhard et al^4^. Following Engelhard, we define the relative contribution for each group of regressors by the reduction in variance explained when setting their betas to zero (referred to in Engelhard et al^4^ as the “no refitting” method), then taking the ratio of these relative to the sum over all features.

Clustering was performed using MATLAB’s ‘fitgmdist’ function on the matrix of neuron x feature contributions, using 1,000 maximum iterations, 0.30 regularization value, 100 replicates, and the covariance matrix constrained to diagonal. To select the number of clusters, we used the fitgmdist function to calculate the AIC score for models with varying numbers of clusters and chose the model with the lowest AIC score. The AIC score served as a measure of model fit given the number of parameters, calculated by taking the difference between 2 times the number of parameters and 2 times the log likelihood of the model. After taking the cluster with the lowest AIC scores, we ordered the relative contribution by those clusters.

### Wall-Texture Analysis

In **Fig. 3**, we identified a repeating wall-texture pattern in the maze by analyzing video frames of a maze with the view angle fixed at 0 degrees (N = 5000 trials). We calculated the similarity matrix for the video frames; specifically, given the video frames, we flattened the video frame at each timepoint into vectors, mean-corrected and normalized the vectors, and measured similarity for all pairs of frames as the cosine of the angle between these vectors (concretely, the i-jth entry of the similarity matrix gives the cosine of the angle between the video frames at times i and j). We also visualized the average, over positions, of each frame’s similarity to those ahead and behind it, as a function of distance. We repeated the same analyses on the feature-specific RPEs, calculating the similarity in the feature-specific RPEs at each timepoint. The feature-specific RPEs here were derived by running the agent with the trained weights from the normal maze described above, but not allowing the agent to change its view angle in the stem of the maze (always fixed at 0 degrees). For **Extended Data Fig. 4**, we repeated the same analysis but calculated the similarity with the position lagged scalar RPE instead.

### Confirmatory versus disconfirmatory cue responses

We defined confirmatory cues as cues that appeared on the side with more evidence so far, and disconfirmatory cues as cues that appeared on the side with less evidence so far. If the agent or mouse had seen an equal number of cues on both sides, the next cue was defined as a neutral cue. For the neural data, we isolated cue kernels as in Engelhard et al^4^ with some modifications: instead of using contralateral and ipsilateral cues, we used predictors including contralateral and ipsilateral cues with contralateral evidence, neutral evidence, and ipsilateral evidence so far. For **Fig. 4b**, we selected those feature-specific RPE units that were modulated by cue onset and the 10 timesteps after cue onset, regardless of left or right cues. For **Fig. 4c**, we selected among the neurons modulated by cues from the encoding model (N = 77/303), plotting only the units modulated by cues (N = 62/303).

For **Extended Data Fig. 5a-b**, we applied the same regression for **Fig. 4c** to the original data split to recordings from the left and right hemispheres. For the feature-specific RPE model responses in **Extended Data Fig. 5c-d**, we split the units to left-versus right-cue responsive units based on the heatmaps in **Fig. 2g**. The units were sorted by eye, choosing the units that responded to the left and right cues respectively at cue onset and throughout the 10 timesteps after cue onset and excluding units that responded at cue offset.

### Outcome Period Responses

In **Fig. 5a,d**, the scalar RPE was calculated by summing the feature-specific RPE units. For the model responses at outcome time for **Fig. 5b,e**, we normalized each unit’s activity 5 timesteps before and 5 timesteps after reward across 5000 trials so the minimum is 0 and the maximum is 1. Then, we averaged across trials and took the response at reward time. For the neural responses at outcome time for **Fig. 5c,f**, we matched the original empirical paper^4^ and calculated the average activity in the first 2 seconds after the onset of the outcome period, baseline corrected by subtracting the average activity from the 1 second period preceding the outcome. For the histograms in **Fig. 5b-c**, **e-f,** a two-sided Wilcoxon signed rank test was performed to determine the p value for the median (yellow line).

### Successor Representation Modeling

We modeled a family of DA responses using a vector of prediction errors for a successor representation, based on Gardner et al^39^. For comparability with the feature-specific RPE model, we used as targets for these successor predictions the same network-derived features 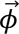 from the feature specific RPE model (LSTM units from **Fig. 2b**). Thus, we have a successor feature model, whose weights we learned using Algorithm 3 of Barreto et al^121^. Given the input state 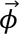 (*s*_*t*_), we trained a set of weights *w* to learn a vector of successor features 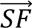 with each *SF*_*i*_ trained with its own sensory prediction error *δ*_*i*_ (**Fig. 6a**). We z-scored the target features 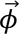 (*s*_*t*_) before training the successor feature model. To calculate the sensory prediction errors (to match with the feature-specific PEs in **Fig. 1c**), we trained a weight matrix *w* with rows of 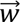 _*i*_, such that each successor feature

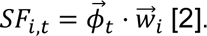

Each 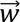 _*i*_ for each successor feature *SF*_*i*_ was then updated using the classic scalar RPE recursive step; that is, each has its own sensory prediction error *δ*_*i*_

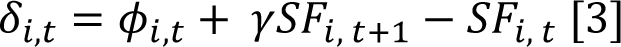

We updated each 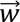 _*i*_ accordingly using:

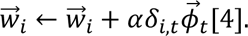

A total of 64 successor features were trained to match with the vector of 64 input features 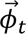. We trained using this recursive update for 15000 trials until the weights converged, using discount factor *γ* = 0.99 and a learning rate *α* = 0.001.

After training our weight matrix *w*, we reran the algorithm with the learned weights frozen to calculate the 64 sensory prediction errors for the 64 features. In **Fig. 6b-d**, we generated the heatmaps with the same methods as with feature-specific RPE model’s results in **Fig. 2e-g**. In our analyses in **Fig. 6e**, we compare the feature-specific RPE model’s scalar RPE with the averaged sensory PE, which is the average over the 64 sensory PEs. The outcome period plots **Fig. 6f-i** was generated the same way as **Fig. 5a-b,d-e**.

### Distributional Reinforcement Learning Modeling

To model the features with a distributional reinforcement learning model, we adapted a TD ™1 algorithm to train weights for 31 distribution channels, based on algorithms specified in refs.^33,122^ and briefly summarized below. Each channel has two learning rates, *α*_*i*_^+^ for positive prediction errors and *α*_i_ ^−^for negative prediction errors, chosen so that the expectiles 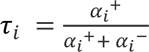 evenly tile the range between 0 and 1 (exclusive) with *α*_*i*_^+^ + *α*_*i*_^−^ = 0.001.

The input features for the model were (again, for comparability with the feature-specific RPE) that model’s network-derived features 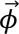, z-scored. We trained weights that mapped the features to distributional value channels, including a bias term. We trained a set of linear feature weights for each channel by at each timestep directly predicting the (*γ*= 0.99 discounted) final outcome of the trial (effectively, expectile regression or TD-1; this avoids distributional imputation in a bootstrapped TD-0 update).

After training for 15000 trials, we froze the converged weights to calculate the 31 expectile prediction errors for the distribution channels. We then computed TD-0 Bellman errors at each step with an imputation algorithm^122^ to sample the one-step bootstrap backup value for each of the 31 channels from the ensemble. Each unit’s per trial positive and negative prediction errors were weighted by its *α*_*i*_^+^ and *α*_*i*_^−^, respectively.

To generate plots for **Fig. 7b-i**, we repeated the same methods as in **Fig. 6b-i**, with one difference for the heatmaps in **Fig. 7b-d**; specifically, in the heatmaps, we used the original ordering of the distribution channels, since the channels themselves already have a natural order from pessimistic to optimistic channels.

### Action Prediction Error Models

The Action Prediction Error (APE) framework may be formally thought of as the standard RL setup but execution of actions provides the reward. Though previous work^40,41^ implemented APE in the manner of the Rescorla-Wagner rule (i.e., without a future-predictive term), we extended these models to a full TD formulation analogous to Schultz et al.^3^ (It is worth noting that Rescorla-Wagner is a special case of this with maximal temporal discounting).

The scalar APE model used to describe the Parker et al.^7^ results correspondingly used a variant of the typical Q-learning approach. For a hypothetical DA population with some preferred action *a* (e.g., left lever press), that is:

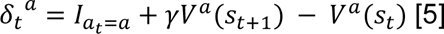

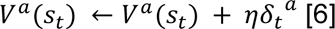

Here, *δ*_*t*_^*a*^ is the prediction error at time *t* and corresponds to the dopaminergic response. *I*_*a* =*a*_ is an indicator for the population’s preferred action: equal to 1 if *a*_*t*_ = *a* and 0 otherwise. *γ* ∈ [0, 1] is a temporal discounting factor. *V*^*a*^(*s*) denotes the value of state *s*, which is equal to the expected cumulative discounted number of times *a* will be executed during the remainder of the trial. *η* is a learning rate parameter.

The feature-based APE model used to describe Jin and Costa^5^ is similar. Indeed, the approach used to derive it was identical; we treated the actions as being the source of the rewards:

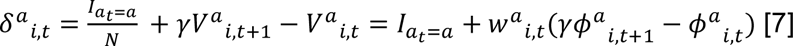

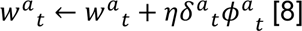

Here, *δ*^*a*^_*i*,*t*_ is the prediction error at time *t* for feature channel *i* and corresponds to the dopaminergic response of that DA neuron. *N* is the total number of channels. *W*^*a*^_*i*,*t*_ is the *i*th entry in the value weight vector at time *t*. *ϕ*^*a*^_*i*,*t*_ is the *i*th entry in the state vector at time *t*.

Our models of the Parker et al^7^ task and the Jin and Costa^5^ task were essentially the same with some slight differences in parameters, so we will first describe their shared abstract structure and then elaborate separately on the parameters. Both tasks began at a “start” state in which levers were presented. In order to randomize the timing of subsequent actions, the start state had a probabilistic self-transition. After exiting the start state, the agent needed to press a lever a fixed number times in order to progress to the outcome states; they chose which lever to press at each timepoint according to a softmax policy over the true reward values for each lever. For each press, they first progressed to a “premotor” state that preceded the chosen lever press state (i.e., led to it with probability 1 on the next timestep) and then to the “press” state itself (where the agent that preferred the corresponding action earned a reward of 1). The final choice state similarly had a probabilistic self-transition, after which the agent entered a reward state or an omission state depending on the levers it had pulled. An abstract state diagram may be found in **Extended Data Fig. 7**.

In the Parker et al^7^ task, the agent only needs to press a lever once to reach the outcome states. The softmax policy has an inverse temperature T = 0.25. The start state self-transition has probability 0.8. The pre-outcome self-transition has probability 0.95. The temporal discount factor for the reward model is *γ*_*RPE*_ = 0.95. The temporal discount factor for the APE model is *γ*_*APE*_ = 0.5. Discounting is harsher for the APE model to match the value used in modeling the Jin and Costa^5^ task (described below) – but we note that the specific values used do not change the qualitative pattern of the results but only affect the scale of the responses in **Fig. 8d, left**.

In our simplification of the Jin and Costa^5^ task, the agent needs to press a lever three times to reach the outcome states. To randomize the within-sequence press timing, each of the choice states also has a self-transition with probability 0.1. The start state self-transition has probability 0.8. The softmax policy has an inverse temperature *τ* = 0.25. The temporal discount factor for both the scalar and feature-specific models is *γ* = 0.5. In general, we believe that harsher discounting is appropriate for the APE models since the computational goal of action prediction is presumably preparing a single upcoming response rather than computing a long-run average– but we note, as above, that the specific value of *γ* does not affect the qualitative pattern of the results (i.e., that there are DA neurons with specific preferred presses that emit a biphasic response, and that those neurons will change their tuning in response to changes in tuning upstream), only the relative magnitudes of the peaks and troughs in the DA response.

### Retraining Network Results

The retrained network results in **Extended Data Fig. 1** used the same hyperparameters as the original run and took 162 million timesteps to learn. One minor difference was the scheduling of the hyperparameter *n*_*steps*_, which we lowered to 20 early in training, then increased to 50 for the last 28 million timesteps. The lower *n*_*steps*_ helped improve performance in part because it meant the A2C algorithm had fewer trajectories it needed to sample and explore and therefore could more easily learn the task. After performance was above chance, a higher *n*_*steps*_ was needed to match the original performance in order to allow the agent to integrate the cues across the entire trial to maintain the correct tower difference count. The plots for **Extended Data Fig. 1** were generated the same as their counterparts in **Fig. 2c-d**, **Fig. 4b**, **Fig. 2e-g**, and **Fig. 5a-b,d-e.**

### Code Availability

Code used for the deep reinforcement learning model, virtual reality environment, and analysis of the data to reproduce the figures can be found at https://github.com/ndawlab/vectorRPE/.

### Data Availability

The data that support the findings of this study will be available on figshare upon publication.

## Notes

### Competing Interest Statement

The authors have declared no competing interest.

### Summary of Updates

Edited and refined some paragraphs for the discussion.

